# Sarm1 Gates the Transition from Protective to Repair Schwann Cell States Following Nerve Injury

**DOI:** 10.1101/2024.07.11.603162

**Authors:** E. Stepanova, S. Hunter-Chang, J. Lee, A. Tripathi, C. Pavelec, C Cho, T. Vegiraju, J. Guo, C. Kim-Aun, S. Kucenas, N. Leitinger, J. Coutinho-Budd, J. Campbell, C. Deppmann

**Affiliations:** Department of Biology, College of Arts and Sciences, Charlottesville, VA 22902, United States of America; Neuroscience Graduate Program, School of Medicine, University of Virginia, Charlottesville, VA 229022, United States of America; Department of Biomedical Engineering, School of Engineering, University of Virginia, Charlottesville, VA 22902, United States of America; Department of Neuroscience, School of Medicine, University of Virginia, Charlottesville, VA 229022, United States of America; Department of Cell Biology, School of Medicine, University of Virginia, Charlottesville, VA 229022, United States of America; Program in Fundamental Neuroscience, College of Arts and Sciences, Charlottesville, VA 22902, United States of America; Department of Pharmacology, School of Medicine, University of Virginia, Charlottesville, VA 229022, United States of America; Robert M. Berne Cardiovascular Research Center, School of Medicine, University of Virginia, Charlottesville, VA 22902, United States of America

## Abstract

Schwann cells (SCs) transition into a Repair state after peripheral nerve injury; however, the early SC injury response preceding this transition remains poorly understood. We demonstrate that Sarm1, a key regulator of axon degeneration, is expressed and upregulated in SCs after nerve injury. Cell-type-specific Sarm1 knockout SCs exhibit enhanced axon protection *in vitro*, and SC- and glia-specific Sarm1 deletion confers axon protection in mouse sciatic nerve and Drosophila wing injury models. Single-nucleus RNA sequencing revealed that Sarm1-deficient SCs are enriched in a distinct cluster expressing genes with developmental roles in axon and myelin protection, with increased oxidative phosphorylation gene expression across all injured SC states. We propose that Sarm1 gates the transition from a Protection-Associated Schwann Cell (PASC) state to a Repair SC state, establishing Sarm1 as a multi-functional regulator with implications for peripheral neuropathies and neurodegenerative diseases.

## Introduction

Schwann cells (SCs), one of the most numerous cell types of the peripheral nervous system (PNS), play a crucial role in supporting axons and promoting regeneration after injury(*1–4*). Following nerve injury, SCs undergo significant changes, transitioning to a repair phenotype characterized by the upregulation of regeneration-associated genes, activation of myelinophagy, and formation of bands of Büngner(*1*, *2*, *5*). This repair phenotype is driven by the activation of transcription factors, such as c-Jun, which is essential for initiating the SC injury response(*6*, *7*).

While the repair SC phenotype is critical for regeneration, this cell state takes time to emerge in mice post-injury(*1*). Recent work demonstrated that within 24-36 hours following injury, SCs undergo a dramatic glycolytic shift driven by mTORC1 signaling and associated transcription factors Hif1α and c-Myc, which enhances bioenergetic support to injured axons through increased lactate production(*8*). This glycolytic response is crucial for delaying axonal degeneration and maintaining axonal integrity during the early injury period.

What happens in SCs during the immediate hours after injury, before this glycolytic shift occurs? Do SCs transition directly from a myelinating/non-myelinating to the glycolytic and a repair state, or do they pass through intermediate transcriptional states? Furthermore, do these putative early-responding SCs serve a distinct functional role in regulating axonal degeneration, separate from their later functions in promoting regeneration? Understanding both the cellular states and functional properties of SCs in these first hours after injury, as well as the molecular mechanisms that regulate transitions between SC states, could lead to targeted therapies that prevent axonal degeneration(*9*, *10*) while uncovering new strategies for enhancing the efficiency of nerve regeneration and promoting better functional recovery(*11*, *12*).

Sarm1, a toll-like receptor adaptor protein, has emerged as a central regulator of axonal degeneration(*13*). NMN accumulates in injured axons following loss of the NAD+ synthesizing enzyme NMNAT2, activating Sarm1(*14*). Activated Sarm1 cleaves NAD+, leading to a rapid depletion of ATP and subsequent axonal degeneration(*15*). Genetic deletion of Sarm1 confers robust axon and myelin protection after injury (*13*, *16*, *17*). Recent studies demonstrate that Sarm1 is also required for the timely induction of the SC repair response following peripheral nerve injury. In the absence of Sarm1, there is a significant delay in the upregulation of c-Jun in SCs, which is essential for initiating their transformation into a repair phenotype(*6*). This is consistent with the finding that Sarm1 loss prevents pJNK induction in the sciatic nerve 30 minutes after injury(*16*). This delay in SC repair program activation creates a unique opportunity: Sarm1 deficiency effectively “locks” SCs in an early post-injury state that precedes the canonical repair response, allowing identification and characterization of this previously elusive early-responding SC population. However, a SC-autonomous role for Sarm1 has not been reported.

Here we investigate how Sarm1 in SCs regulates axon protection following nerve injury. Using *in vitro* injury models and *in vivo* mouse experiments, we demonstrate that SC Sarm1 deletion confers axon protection, confirming that this effect is SC-autonomous. Furthermore, Sarm1 knockdown in Drosophila glia recapitulates this protection, indicating that glial-mediated axon preservation is evolutionarily conserved and is independent of myelination status. To define the underlying transcriptomic basis of this protection, we leveraged Sarm1 knockout mice and single-nucleus RNA sequencing at multiple timepoints after nerve injury. We propose that SCs adopt a Protection Associated Schwann Cell (PASC) state immediately following injury—a transient phenotype that precedes the canonical repair program and is characterized by an intermediate pseudotime position between homeostatic (myelinating and non-myelinating SCs) and Repair states, expression of myelin formation and axon protection genes (Fth1, Ptgds, Ptprd, Trf). Our findings reveal that Sarm1 gates the transition from the PASC state to the repair SC, with Sarm1-deficient SCs enriched in the PASC state and exhibiting upregulation of mitochondrial respiration genes, though whether this metabolic profile defines PASCs in wild-type cells or reflects a specific consequence of Sarm1 loss requires further investigation. This work establishes PASCs as a distinct functional state preceding the canonical repair program, providing a window of opportunity for axonal preservation before degeneration occurs, with broad implications for treating peripheral nerve injuries and neurodegenerative diseases.

## Results

### Sarm1 mRNA expression in Sox10 positive cells

Recent findings implicate Sarm1 in SC function beyond its established axon-autonomous role: Sarm1 is crucial for the timely induction of the SC repair response following peripheral nerve injury(*18*, *19*). Moreover, the delay of Sarm1KO entry into a Repair SC state creates an opportunity to study a novel, pre-Repair SC cell state in Sarm1KO animals. Previous work in zebrafish detected low-level Sarm1 mRNA expression in approximately 20% of SCs on the posterior lateral line nerve (PLLn)(*20*).

We examined this by performing Sarm1 and Sox10 RNA fluorescence in situ hybridization (RNA FISH) on WT sciatic nerves (Figure 1C-E; S1B). Consistent with the zebrafish findings, we detected 19% of Sox10+ nuclei expressing Sarm1 (Figure S1A), with an average of 0.25 Sarm1 puncta per Sox10+ cell (Figure S1A). When we extended the nuclear boundary to approximate cell membrane boundaries, the number rose to 0.73 transcripts per cell and 65% of Sox10+ cells expressing Sarm1 (Figure 1D,E). 3D reconstruction using IMARIS software confirmed Sarm1 RNA puncta localized within SCs (Figure S1 C-D).We observed no differences in the number of Sox10+ cells expressing Sarm1 after injury, but detected an increase in the number of transcripts per nucleus and per cell (Figure 1C-E, S1A). Since increased Sarm1 transcription requires intact nuclei with active transcriptional machinery, and after injury neuronal cell bodies (and their nuclei) are disconnected from distal axons, the observed Sarm1 transcripts are highly unlikely to be of axonal origin. We further confirmed SC-autonomous Sarm1 expression by detecting Sarm1 mRNA in cultured SCs from WT mice, which was absent in cultures derived from Sarm1KO mice (Figure 1A, B; S1E). This observation is consistent with previous work identifying a significant increase in Sarm1 transcripts in WT cultured SCs compared to Sarm1KO counterparts(*20*).

**Figure 1:**
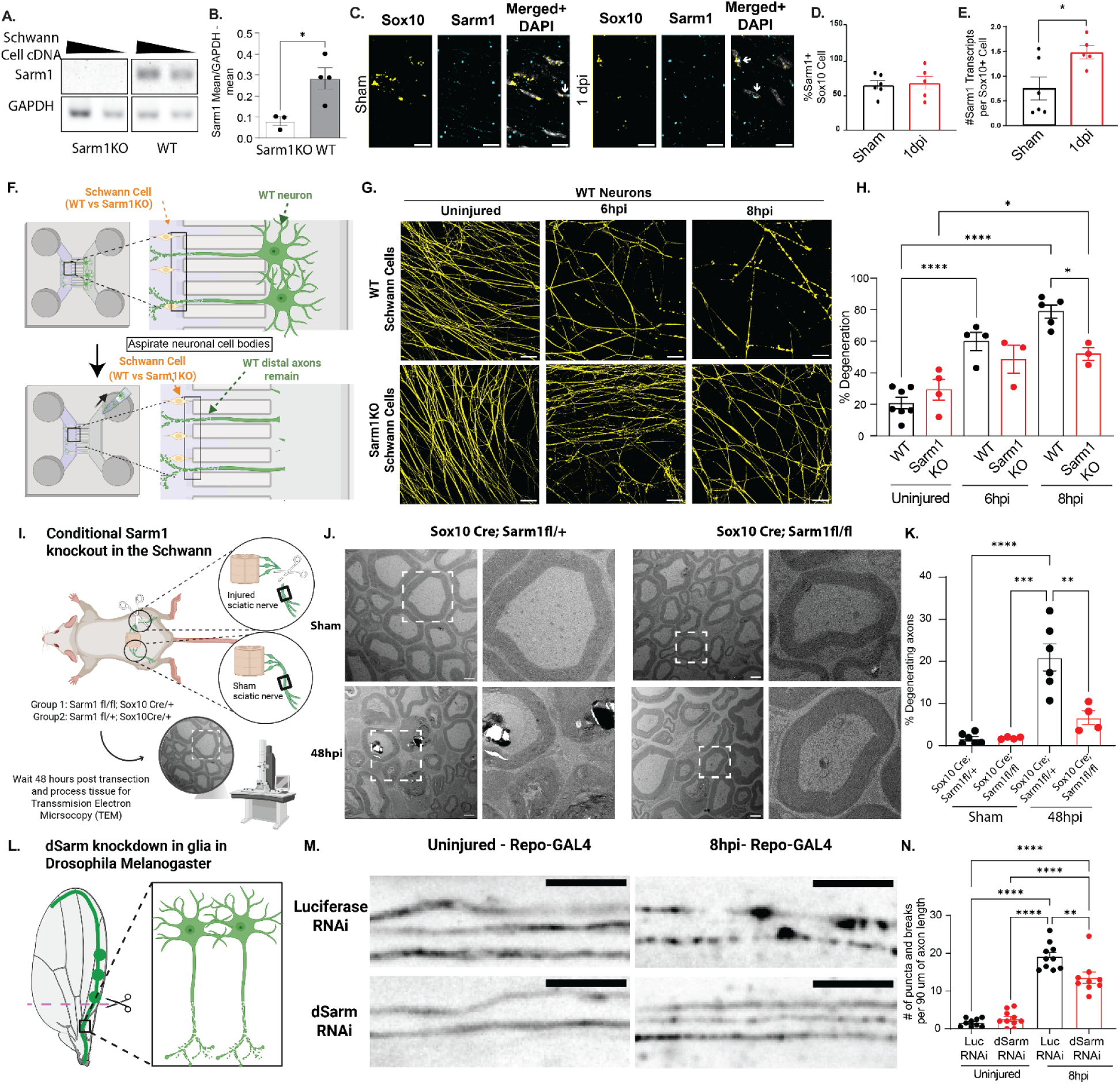
Sarm1/dSarm’s Role in Axon Degeneration in Schwann Cells. **A-B.** Sarm1 expression in primary cultured SCs. SCs were isolated from mouse sciatic nerves at P0-P2 and maintained *in vitro* for one week. RNA was reverse transcribed into cDNA and primers for Sarm1 and GAPDH were used to amplify the respective transcripts. **A.** RT-PCR analysis showing Sarm1 and GAPDH transcript levels. Two cDNA concentrations (1× and 0.5×) were loaded per sample, as indicated by triangles above the gel images. **B.** Quantification of Sarm1 expression normalized to GAPDH. Band intensities were measured from the gel in (A) and normalized to corresponding GAPDH bands. Values represent the average of both cDNA concentrations for each biological replicate. n = 3 independent litters (Sarm1KO) and n = 4 independent litters (WT). Data presented as mean ± SEM. Statistical comparison by Welch’s t-test. **C.** Representative images of RNA in situ hybridization targeting Sox10 (yellow) and Sarm1 (cyan) transcripts in sham and 1 day post-injury (dpi) sciatic nerves. Merged images include DAPI (gray). Scale bar = 10 µm. Each data point represents one mouse (n = 6 WT, n = 5 Sarm1KO), calculated as the mean of 2-7 sections per mouse. **D.** Quantification of the percentage of Sox10+ nuclei co-expressing Sarm1. Data presented as mean ± SEM. **E.** Average number of Sarm1 transcripts per Sox10+ nucleus. Data presented as mean ± SEM. **F**. Schematic representation of a co-culture system with wild-type (WT) and Sarm1 KO SCs cultured with wild-type neurons, followed by aspiration axotomy. Boxed area indicates region of imaging. **G.** Representative images of Superior Cervical Ganglia co-cultured with either WT or Sarm1KO SCs, under uninjured conditions 6 and 8 hours post-injury (hpi). Axons visualized by Tuj1 antibody staining. Scale bar = 50 µm. **H**. Quantification of axon degeneration from images in (**G**), performed by an investigator blinded to experimental conditions. Each dot represents a percent degenerating axons (∼100 -300 axons quantified) per animal. Two way ANOVA with Tukey’s test for multiple comparisons.p-value**** <0.0001, *** 0.0002 **0.0021. Data presented as mean ± SEM. **I**. Graphic representation of sciatic nerve injury in mouse sciatic nerve followed by Transmission Electron Microscopy (TEM). **J**. Representative images from TEM showing conditional Sarm1 knockout in Schwann cells (Sox10 Cre; Sarm1fl/fl) vs controls (Sox10 Cre; Sarm1fl/+) Sham vs 48hours post injury (48hi). Scale bar=2um **K.** Quantification showing % of degenerating axons compared to the total number of large caliber axons quantified for a given animal. Each dot represents one animal with a minimum of 100 axons quantified. Two way ANOVA with Tukey’s test for multiple comparisons, p-value ****<0.0001, **0.0050. Data presented as mean ± SEM. **L.** Graphic representation of wing injury in Drosophila melanogaster. A unilateral lesion in the wing was achieved by ablating all visible cell bodies. 8 hours after injury, the injured wing and contralateral uninjured wing was collected for immediate imaging. **M**. Representative image of control (Luciferase RNAi) and experimental (dSarm RNAi) Vglut-QF2 MARCM labelled CD8::GFP+ neurons crossed to the pan-glial RepoGal4 driver. Scale bar is 10um. **N.** Blinded investigator quantified the number of puncta and breaks per 90 micron of axons length as a proxy for axon breakdown. Data reported as mean +/- SEM. Two way ANOVA with Tukey’s test for multiple comparisons, p-value ****<0.0001, **0.0050.

Importantly, Sarm1 protein expression has been reported in oligodendrocytes but not SCs(*20*). The apparent discrepancy between RNA and protein levels may be explained by the differences in detection sensitivity between in situ hybridization and immunohistochemistry, along with the poor signal-to-noise ratio of the Sarm1 antibodies we have evaluated.

### Sarm1KO SCs confer improved protection against WT axon degeneration in an *in vitro* injury model

Having established that Sarm1 mRNA is expressed in SCs and upregulated following injury, we next asked whether this SC-intrinsic Sarm1 expression contributes functionally to axon degeneration. Sarm1’s role as a central executioner of Wallerian degeneration is well established, and loss of neuronal Sarm1 is sufficient to robustly protect axons after injury (*13*). However, sufficiency does not preclude additional non-cell-autonomous contributions from neighboring cells — a possibility that has received little direct experimental attention. To test this, we established an *in vitro* microfluidic co-culture system (Figure 1F) that allows for the spatial separation of neuronal cell bodies and distal axons(*21*). WT superior cervical ganglion (SCG) neurons were cultured and allowed to extend axons to the distal chamber of the microfluidic device. WT or Sarm1KO SCs were then added to the axon chamber. Following axotomy, achieved by removing the neuronal cell bodies, we assessed axon degeneration in WT SCG neurons co-cultured with each SC genotype.

WT SCG neurons co-cultured with WT SCs showed significant axon degeneration at both 6 and 8 hours post-injury (Figure 1G, H). In contrast, neurons co-cultured with Sarm1KO SCs did not exhibit significant degeneration until 8 hours post-injury, representing a significant 26.8% reduction in axon degeneration compared to the WT SC condition. Together, these results demonstrate that Sarm1 expression in SCs contributes to injury-induced axon degeneration *in vitro*, and that its loss enhances the capacity of SCs to protect axons from degeneration.

### Enhanced Axon Protection Following Sciatic Nerve Injury in Mice with Schwann Cell-Specific Sarm1 Deletion

To investigate whether Sarm1 plays a role in mammalian SCs *in vivo*, we conditionally knocked out Sarm1 in SCs using Cre-lox technology. Mice expressing Sox10-Cre were crossed with Sarm1 flox mice to generate Sox10 Cre/+; Sarm1fl/+ controls and Sox10 Cre/+; Sarm1fl/fl conditional knockouts. Both genotypes underwent sciatic nerve transection surgery, and nerves were collected for transmission electron microscopy (TEM) to assess axon and myelin morphology following injury (Figure 1I)

In sham nerves, the proportion of degenerating axons was minimal and comparable between genotypes (1.7% and 1.8% for controls and conditional knockouts, respectively) (Figure 1J-K). At 48 hours post-injury, control mice showed a significant increase in axon degeneration, reaching 21%. In contrast, SC-specific Sarm1 conditional knockout mice exhibited substantially less degeneration, with only 6.7% of axons degenerating at the same timepoint. These findings demonstrate that SC-expressed Sarm1 promotes axon degeneration *in vivo*, consistent with our *in vitro* observations.

Taken together, these results reveal a previously unrecognized role for Sarm1 in SCs, demonstrating that Sarm1-driven axon degeneration can occur non-cell-autonomously during the early stages of nerve injury.

### Glial dSarm knockdown provides improved protection against axon degeneration in an *in vivo Drosophila melanogaster* wing injury model

To investigate whether the glial role of Sarm1 in axon degeneration is evolutionarily conserved, we turned to *Drosophila melanogaster*. This model offers powerful cell-type-specific genetic tools, functional conservation of glial roles, and evolutionary conservation of dSarm’s role in axon degeneration(*13*, *18*). Importantly, Drosophila wrapping glia are functionally analogous to SCs but lack myelination capacity, providing a convenient system to test whether the glial role of Sarm1 extends to non-myelinating SC-like cells. This is particularly relevant because, while our *in vivo* mouse experiments assessed degeneration of large-caliber myelinating axons, our *in vitro* co-culture system used non-myelinating SCs — leading us to predict that the glial contribution of Sarm1 to axon degeneration operates independently of myelination status.

We first validated two dSarm RNAi lines by crossing them with the neuronal driver Dpr1-Gal4, which labels approximately 40 neurons and their axons with CD8-tagged GFP. Partial wing ablation was used to remove all visible cell bodies (Figure 1L), and axon degeneration was assessed at consistent locations along the marginal wing vein 3 days post-injury, with the contralateral uninjured wing serving as a control. Only one dSarm RNAi line (line #2) demonstrated significant protection relative to Luciferase RNAi controls (Figure S1F,G), and this line was used for all subsequent glial experiments.

To assess dSarm’s role specifically in glia, we crossed UAS-Luciferase RNAi or UAS-dSarm RNAi with the pan-glial RepoGal4 driver, utilizing MARCM labeling to visualize individual axons as previously described. To align with the early degeneration window observed in our mouse models, we assessed axon integrity at 8 hours post-injury rather than 3 days. At this timepoint, control wings exhibited significantly more axonal puncta and breaks than wings with glial dSarm knockdown (Figure 1M,N), demonstrating that glial dSarm accelerates axon degeneration during the early injury response.

Together, these findings confirm that the SC contribution of Sarm1 to axon degeneration is evolutionarily conserved and is not restricted to myelinating cells.

### Single-nucleus RNA sequencing reveals distinct cell populations in WT and Sarm1 knockout mouse sciatic nerves

Based on our findings that Sarm1 mRNA is expressed in mouse SCs and that Sarm1KO SCs confer protection against axon degeneration (Figure 1), we sought to better understand the role of Sarm1 in the SC injury response. Recent work demonstrates that Sarm1 is required for the timely induction of the SC repair program following peripheral nerve injury(*19*), suggesting that Sarm1 deficiency may lock SCs in a pre-repair state that retains a protective capacity against axon degeneration at early stages of injury. To explore how loss of Sarm1 alters SC transcriptional responses, we employed single-nuclei RNA sequencing (snRNA-seq) to profile sciatic nerves from WT and Sarm1KO mice at defined timepoints following injury (Figure 2A). Given that Sarm1 expression was detected in SCs and that glia- and SC-specific Sarm1 knockout is sufficient to confer axon protection, we reasoned that the transcriptional changes observed in Sarm1KO nerves would primarily reflect SC-intrinsic effects. The need to pool large amounts of tissue to achieve sufficient cell capture also made whole-nerve preparations from Sarm1KO mice the most practical approach.

**Figure 2:**
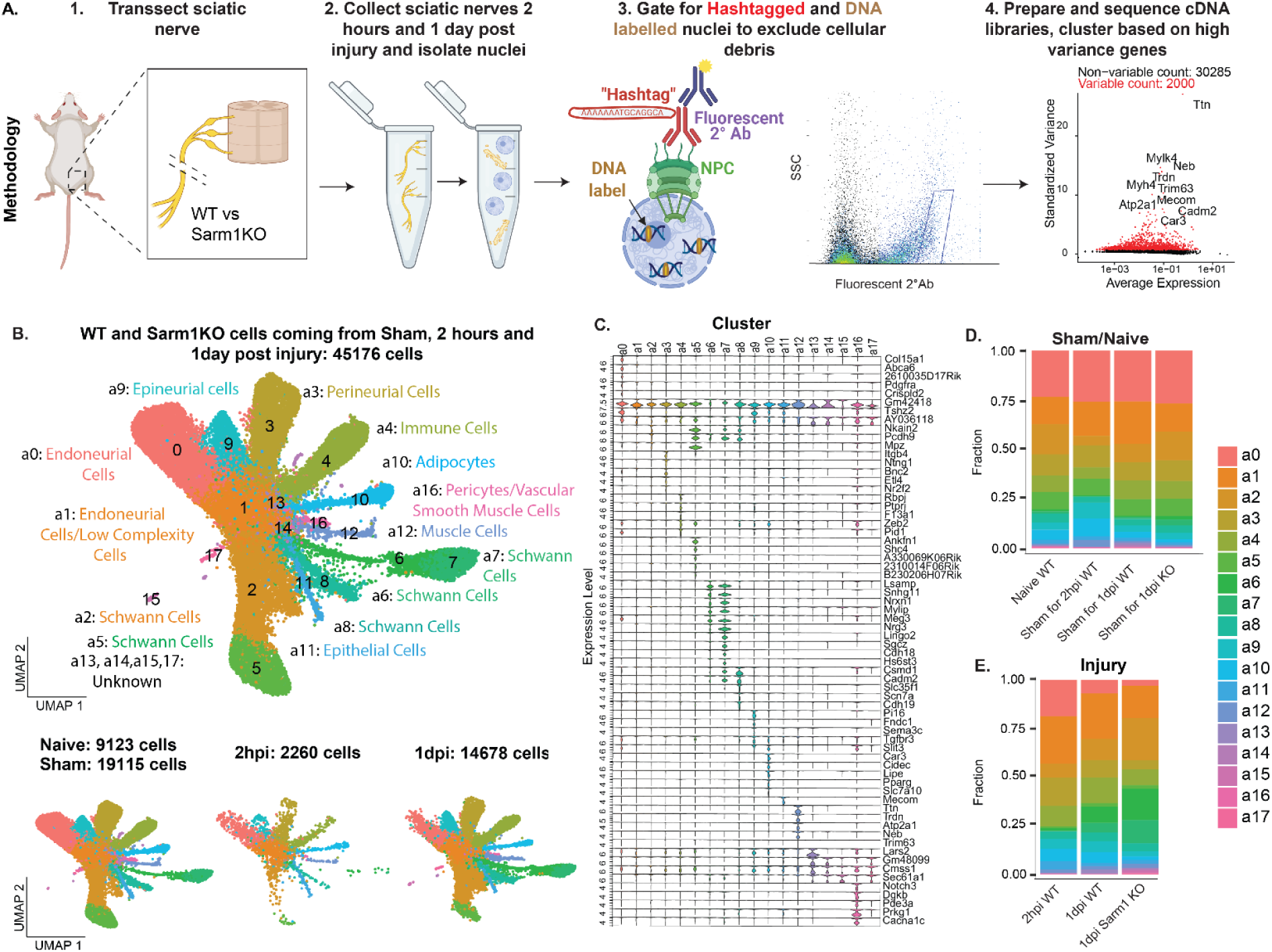
Overview of single-nucleus RNA sequencing (snRNA-seq) in WT and Sarm1KO mouse sciatic nerves. **A.** Schematic workflow of snRNA-seq protocol: (1) surgical transection of right sciatic nerve with left sciatic nerve as sham control; (2) nerve collection 1 day post-surgery and nuclei isolation via Dounce homogenization and density-gradient purification; (3) FACS purification of nuclei using DNA-barcoded antibodies against Nuclear Pore Complex (NPC) and DNA intercalator counterstain; (4) cDNA library preparation, sequencing, and data analysis. **B**. UMAP visualization of single-nucleus transcriptomes. Top: Clustering of nuclei from sham and injured sciatic nerves of WT and Sarm1KO mice (both sexes) at 2 hours and 1 day post-injury. Bottom: Split UMAP plots showing distribution and number of nuclei by condition (sham/naive, 2 hours post-injury, and 1 day post-injury). **C.** Violin plots depicting relative expression levels of marker genes for each identified cluster. **D-E.** Stacked bar plots showing the proportion of nuclei from each cluster by condition for sham (**D**) and injured (**E**) samples.

We surgically transected the right sciatic nerves of male and female 3-6-month-old mice coming from wild type (WT) or Sarm1 knockout (Sarm1KO) backgrounds, while the left nerves served as sham controls (Figure 2A). WT nerves were collected at sham, 2 hours (2hpi), and 1 day (1dpi) post-injury, while Sarm1KO nerves were collected at sham and 1dpi. Since transcriptional changes precede the protein-level and structural changes that determine axon fate, we reasoned that profiling SCs at these timepoints would capture the gene expression programs driving the morphological differences we observe between genotypes at 48 hours. The 2hpi timepoint captures the earliest transcriptional response before major metabolic reprogramming, while 1dpi represents the more elaborated program just prior to the divergence in axon outcomes.

We isolated nuclei from the nerves 1 day post-injury, labeled them for purification by fluorescence-activated cell sorting (FACS) based on nuclei markers (Figure 2A, Figure. S2A), followed by cDNA library preparation using 10x Genomics and sequencing. Clustering 45,176 single-nuclei transcriptomes (∼cells) from WT and Sarm1KO mice based on expression of the top 2000 highest variance genes revealed 17 cell clusters. This included naive, sham, 2hpi and 1dpi experimental conditions (Figure 2B), with 9123 cells coming from naive, 19,115 from sham, 2260 from 2hpi, and 14,678 from 1dpi. We compared the gene expression profiles of the cell clusters to determine their molecular identities. Our results revealed many canonical cell type marker genes enriched in each cluster, allowing us to identify clusters a2, a5, a6, a7 and a8 as SCs based on their expression of genes such as *Mbp*, *Cadm2*, *Ncam1* (Figure S2I). We annotated other clusters as known cell types based on top differentially expressed genes in our analysis (Figure 2C, S2B) and compared to gene profiles reported previously in single-cell RNA sequencing atlases of sciatic nerves(*22*, *23*). Clusters with low cell numbers a13, a14, a15 and a17 could not be confidently annotated due to low gene expression and number of transcripts detected (Figure S2C-E), suggesting these may represent transcriptomes of lower molecular diversity of quality. Based on the gene expression for each cluster (Supplementary Table 1), we annotated cell clusters as the following cell types: endoneurial cells (a0,a1), adipocytes (a10), SCs (a2,a5,a6,a7,a8), perineurial cells (a3), immune cells (a4), epithelial cells (a11), muscle cells (a12), epineurial cells (a9) and vascular smooth muscle cells (a16) (Figure 2B). The proportional distribution of cells within each cluster did not grossly differ across conditions for sham samples (Figure 2D, Figure S2J). For injured samples, overall trends were similar (Figure 2E, Figure S2K), though we observed an increase in the SCs in clusters a6 and a7 at 1dpi compared to 2hpi in WT samples, with an even greater increase in the Sarm1KO 1dpi condition (Figure 2E, S1G,H).

### SCs in uninjured sciatic nerves of WT and Sarm1KO mice exhibit few differences in gene expression

Before comparing sham and injured SCs between genotypes, we analyzed the transcriptional differences between genotypes in sham sciatic nerves. Selective clustering of sham SCs (Figure S3A-C) revealed seven distinct clusters of 4,724 sham SCs: ssc0-ssc6 (Figure S3C) with distinct gene expression profiles for each cluster (Figure S3E, Supplementary Table 3). Cluster ssc4 expressed markers characteristic of non-myelinating SCs, such as *Cadm2* and *Ncam1*(*3*), while clusters ssc0, ssc1, ssc3 and ssc6 were enriched with myelinating SC markers, such *Mbp*, *Mpz* and *Prx* (Figure S3D). Cluster ssc2 is enriched with calcium signaling (*Camk1d*), mitochondrial protein synthesis (*Cmss1* and *Lars2*), and lysosomal degradation of glycosphingolipids (*Hexb*) markers. Cluster ss5 is strongly enriched for neurexins and contactin-associated proteins (*Nrnx1*, *Cntnap2* and *Nrg3*) , which are critical for neuron-glia communication and synapse formation.

WT and Sarm1KO SCs were relatively evenly distributed across these clusters (Figure S3F-H). To assess baseline differences between genotypes, we performed differential expression analysis comparing WT and Sarm1KO sham SCs. Using pseudobulk analysis of 21,991 genes, we failed to detect any genes that differed significantly in expression between the sham genotypes (Figure S3I, Supplementary Table 4). This absence of transcriptional differences suggests that Sarm1 does not regulate SC gene expression in uninjured nerves.

### SCs exhibit distinct transcriptional profiles in WT and Sarm1 knockout sciatic nerves 1 day after injury

To investigate differences between WT and Sarm1KO SCs following 2 hours post injury (2hpi) WT cells and 1 day post injury (WT and Sarm1KO), we selectively re-clustered the SCs from injured nerves (Figure S4A-C). This analysis yielded seven clusters of injured nerve SCs: isc0, isc1, isc2, isc3, isc4, isc5, isc6 (Figure S4C) totalling 7,082 nuclei. The 2hpi WT condition yielded 304 nuclei, 1dpi WT 1,340 nuclei and 1dpi Sarm1KO 5,709 nuclei. Each cluster exhibited a distinct transcriptional profile (Figure S4D and Supplementary Table 5). Top marker genes included *Plp1*, *Pcdh9*, and *Nkain2* for isc0; *Meg3*, *Snhg11*, and *Nrg3* for isc1; *Trpm3*, *Gpc5*, and *Lsamp* for isc2; *Camk1d*, *Lars2*, and *Arhgap24* for isc3; *Cntn5* and *Tead1* for isc4; *Fth1* and *Pcdh9* for isc5 (which shares similarity with isc0); and *Ptgds* for isc6 (Figure S4D,H,I and Supplementary Table 5).

WT 1dpi SC nuclei were abundant in cluster isc3 while Sarm1KO nuclei were more abundant in clusters isc0 and isc1 (Figure S4E-G). Cells from WT 2hpi were predominantly present in cluster isc4, whereas WT 1dpi and Sarm1KO 1dpi cells were more broadly distributed. This uneven distribution suggests that cells from different time points and genotypes occupy distinct transcriptional states along the injury response continuum. We therefore next analyzed the complete dataset including both sham and injured cells to examine the evolution of SC responses to injury.

### Sarm1 deletion enriches for intermediate SC State 1 day after sciatic nerve transection

To determine response of SCs to nerve injury, we combined both sham and injured SCs datasets and reclustered them into eight molecularly-distinct clusters (sc0-sc7; Figure 3A,B). We detected 12,002 total SCs. Clusters sc0 and sc5 both express *Dlc1*, *Lars*, and *Arhgap24*; clusters sc1 and sc6 express *Snhg11*, *Meg3*, *Nrg3* and *Nrg1* (Figure 3D, G, Supplementary Table 2). Clusters sc2 and sc4 are related, both expressing *Fth1*, *Pde4b* and *Ptgds*, though sc4 shows higher expression levels. Cluster sc3 expresses *Prx*, *II16* and *Myo1b*, while cluster sc7 expresses *Csmd1*, *Cdh19* and *Scn7a*, consistent with non-myelinating SCs.

**Figure 3:**
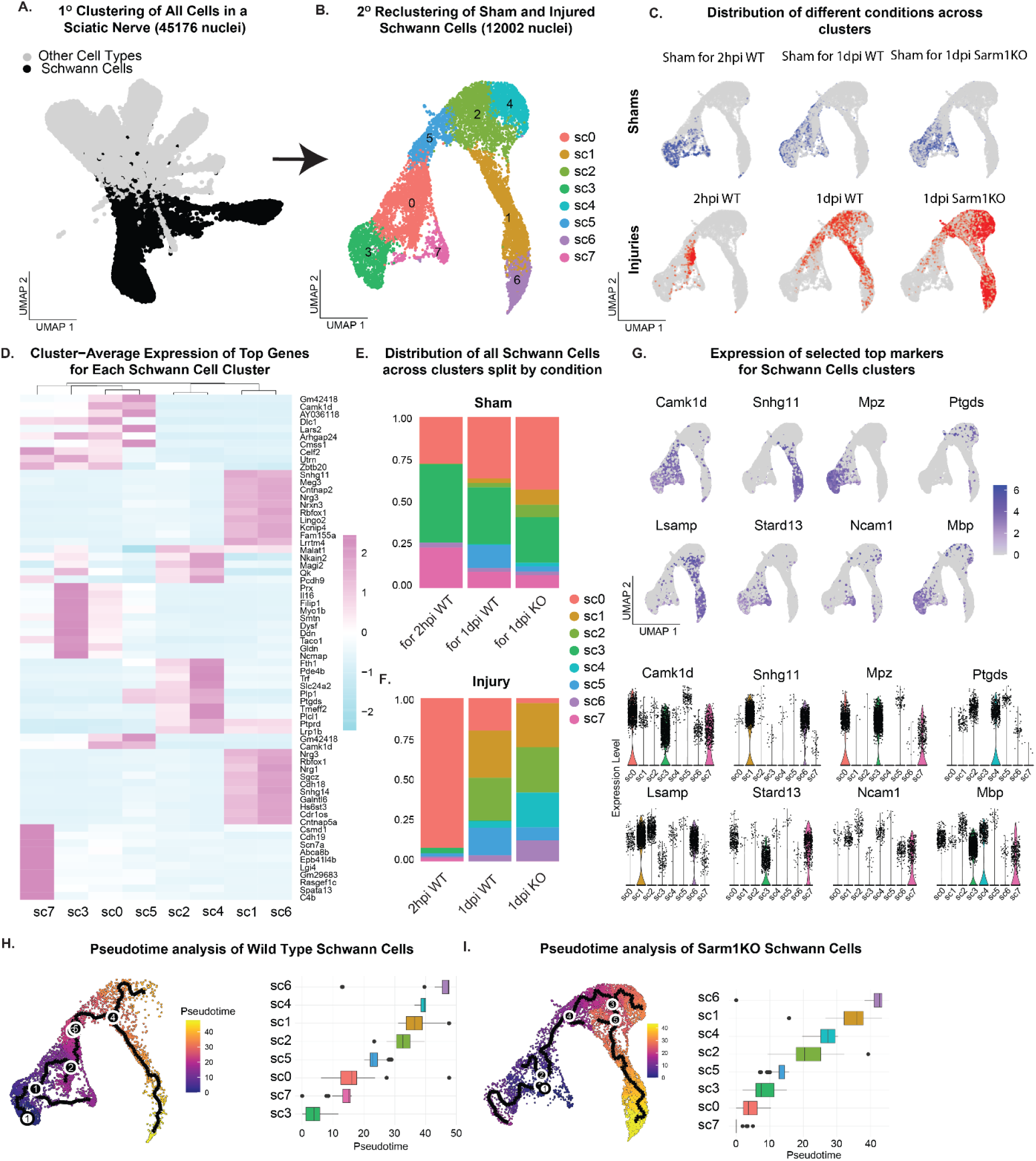
Transcriptional profiling of SCs in WT and Sarm1KO sciatic nerves sham, 2 hours and 1day post injury. **A.** UMAP of full dataset highlighting SC clusters a2, a5, a6, a7, a8 (in black) selected for reclustering analysis. **B.** UMAP of reclustered SCs coming from sham and injured conditions **C.** UMAPs of SCs split by genotype (WT or Sarm1KO) and condition (sham, 2 hours post-injury (2hpi), and 1 day post-injury (1dpi)). **D.** Heatmap showing top 10 differentially expressed genes in each cluster (based on Average Log2FC). **E-F.** Distribution of sham (E) and injured (F) SCs across the clusters for each time and genotype. **G.** Expression of select top markers for each cluster displayed on UMAP overlay and violin plots. **H-I.** Pseudotime analysis of wild type (H) and Sarm1 knockout (Sarm1KO) (I) SCs with sham condition set as time point zero. UMAPs are color-coded by pseudotime values, and bar plots show the pseudotime distribution of each cluster.

Cell distribution across clusters varied by time point and genotype (Figure 3C). Sham SCs predominantly occupied clusters sc0, sc3, and sc7 regardless of the genotype (Figure 3C,E), while 2hpi WT cells were concentrated in cluster sc0 (Figure 3F). At 1dpi, both WT and Sarm1KO cells showed broader distributions with one critical difference: cluster sc4 predominantly comprised Sarm1KO cells with few WT cells. This cluster is enriched for genes essential to myelin formation and axon protection, including *Ptgds*(*24*)*, Ptprd*(*25*), *Trf*(*26*) as well as *Fth1* (Figure 3D, 3G, Supplementary Table2).

Pseudotime analysis with sham cells as the starting point revealed that cluster sc6 represents the most transdifferentiated state in both genotypes (Figure 3H,I). Notably, cluster sc4 appeared more transdifferentiated in the WT trajectory (Figure 3H) but remained less transdifferentiated in the Sarm1 KO trajectory (Figure 3I). This difference suggests that sc4—which is highly enriched in Sarm1 KO cells at 1dpi — represents an intermediate transcriptional state between homeostatic SCs (i.e. closest to the uninjured state) (sc0, sc3, sc7) and repair-like SCs (i.e. most transdifferentiated) (sc1, sc6). Beyond this divergence, WT and Sarm1 KO cells followed similar trajectories, with cells progressing from homeostatic clusters through intermediate states (sc2 and sc5) toward the most transdifferentiated repair-like states (sc1, sc6).

### Sarm1KO SCs upregulate mitochondrial respiration genes 1 day after injury

Our clustering analysis revealed that injury differentially affects gene expression in WT and Sarm1KO SCs. To investigate these differences, we compared the transcriptional responses of each genotype to injury in SCs using a pseudobulk analysis approach (Figure 4A, B, Supplementary Table 6). WT SCs primarily downregulated genes post-injury, while Sarm1KO SCs exhibited more upregulated genes. Top differentially regulated genes in WT SCs included *Ccr5*, *Esrrg*, and *Slc35f4* (upregulated) and *Mpz*, *Sema5a*, and *Mlip* (downregulated) (Figure 4A, Supplementary Supplementary Table 6), suggesting that SCs are mounting an immune response and undergoing demyelination, as evidenced by upregulation of Ccr5 and downregulation of Mpz. In contrast, Sarm1KO SCs showed upregulation of genes including *Rab3c*, *Bex*, and *Syt15*, and downregulation of *Tec*, *Egr2*, and *Lox* (Figure 4B, Supplementary Table 6). Notably, loss of Egr2 (Krox20), a master transcription factor essential for myelination(*27*), suggests that Sarm1KO SCs are responding to injury at the transcriptional level, though through a distinct program from WT SCs.

**Figure 4:**
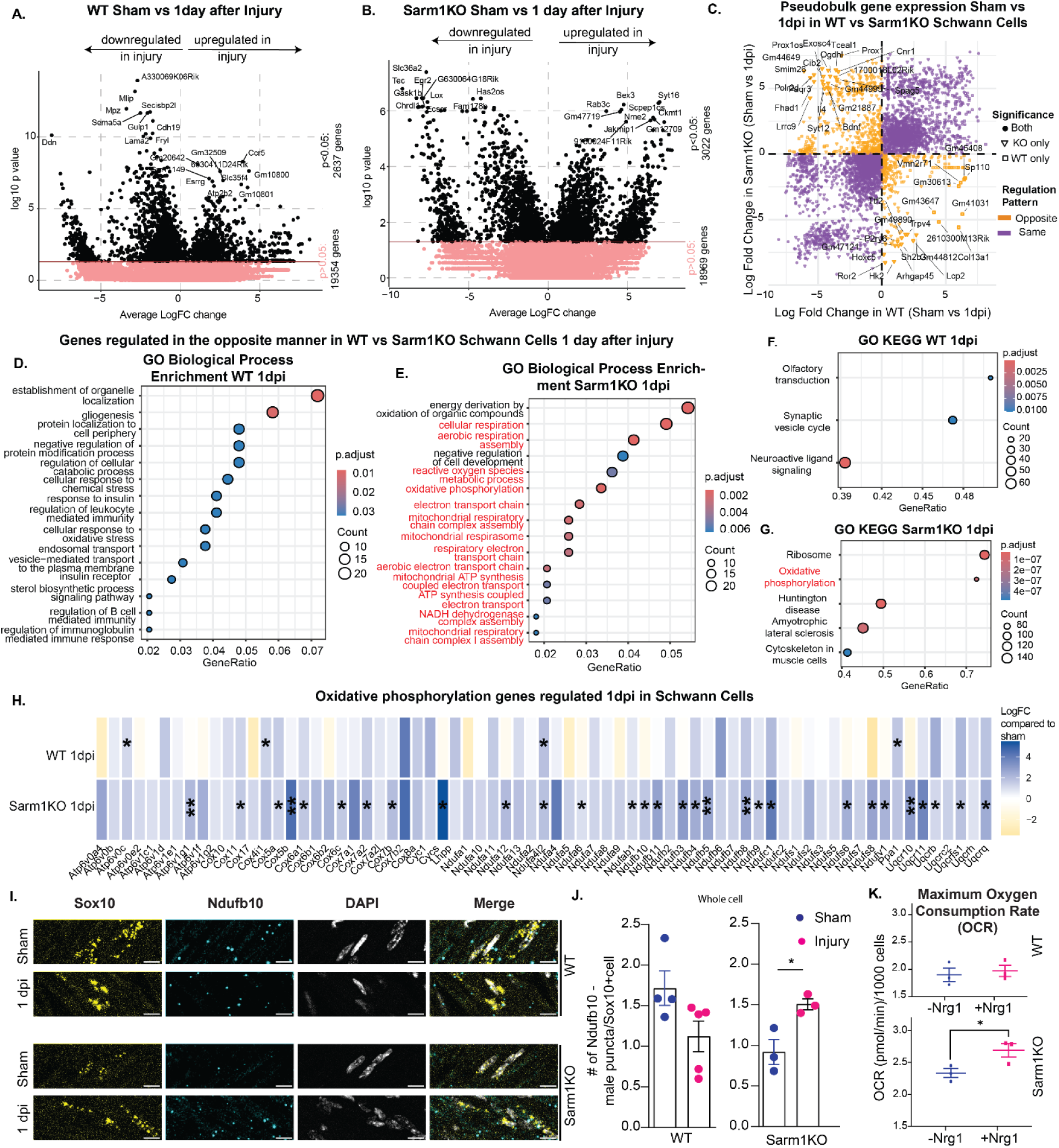
Sarm1KO SCs upregulate mitochondrial respiration one day post-injury. **A-B.** Volcano plots depicting differentially expressed genes 1 day after injury, as revealed by pseudo-bulk analysis in (A) WT and (B) Sarm1KO SCs. **C.** X-Y plot depicting gene expression log fold change 1 day after injury in WT SCs (x-axis) and Sarm1KO SCs (y-axis). Genes regulated in a similar fashion (both upregulated or both downregulated after injury in both genotypes) are highlighted in purple, while genes regulated in opposite directions (e.g. upregulated in one genotype and downregulated in the other) are highlighted in orange. The shape of the marker denotes significance: circles indicate genes significantly regulated in both genotypes according to pseudo-bulk analysis, triangles indicate genes significantly regulated only in Sarm1KO SCs, and squares indicate genes significantly regulated only in WT SCs. **D-E.** Gene Ontology (GO) analysis of biological processes for genes regulated in opposite directions (orange in C) in WT (D) and Sarm1KO (E) SCs. **F-G.** Top enrichment results from KEGG pathway analysis showing expression of all detected genes in WT (F) and Sarm1KO (G) SCs. **H.** Heatmap of mitochondrial respiration genes identified from KEGG analysis (F-G), plotted based on pseudo-bulk analysis, p-value *<0.05, **<0.01, ***<0.001. **I.** Representative images of RNA in situ hybridization for Ndufb10 (mitochondrial respiration gene) and Sox10 (SC marker) in WT and Sarm1KO sciatic nerves under sham conditions and 1 day post-injury. Scale bar = 10 µm. **J.** Quantification of Ndufb10 transcript numbers in Sox10-positive cells from WT and Sarm1KO male sciatic nerves under sham and 1 day post-injury conditions. Data presented as mean ± SEM, p-value * = 0.0256. Each data point represents one mouse (n = 4-5 WT, n = 3 Sarm1KO), calculated as the mean of 3-10 fields of view per mouse. **K.** Maximum oxygen consumption rate (OCR) measurements in primary mouse SCs from WT or Sarm1KO backgrounds using Seahorse XFe96 analyzer. Each data point represents the mean of technical replicates from one mouse litter (3 litters per genotype). Statistical comparison by unpaired two-tailed t-test (± Neuregulin 1), p-value * = 0.0486.

Due to the large number of dynamic genes, we compared genes regulated in opposite directions between WT and Sarm1KO SCs relative to their sham counterparts(Figure 4C). This revealed numerous genes with opposite regulation patterns. For example, *Bdnf*, *Prox1*, and *Fhad1* were upregulated in Sarm1KO but downregulated in WT SCs, while *Trpv4*, *Col13a1*, and *Vmn2r71* showed the reverse pattern. To gain biological insight based on these differences in gene expression, we performed Gene Ontology (GO) analysis on the oppositely regulated genes.

One of the most striking differences between WT and Sarm1KO SCs 1 day after the injury is their metabolic signature. While WT SCs activate programs involving structural remodeling and signaling (Figure 4D), Sarm1KO SCs distinctly upregulate oxidative phosphorylation pathways (Figure 4E). This metabolic shift appears across multiple layers of GO analysis—from Biological Processes (Figure 4D,E), Cellular Processes (Figure S6C,D) to the KEGG pathway analysis (Figure 4F,G). Since oxidative phosphorylation emerged as the most enriched pathway across multiple GO analyses in Sarm1KO cells, we examined the specific genes in this KEGG pathway, plotting their expression as log-fold change compared to sham controls (Figure 4H).

Our single-cell and pseudobulk analysis revealed upregulation of oxidative phosphorylation genes in Sarm1KO SCs with injury, (Figure 4H, Supplementary Table 6) but not in WT SCs. Specific mitochondrial respiration genes—including *Ndufb10*, *Ndufb5*, *Cox5b*, and *Uqcr10*—were significantly upregulated in Sarm1KO SCs in response to injury (Figure 4H). To validate these findings, we performed RNA FISH for *Ndufb5* and *Ndufb10* and observed similar trends (Figure 4I,J, S5D,E). *Ndufb10* expression was significantly upregulated in Sarm1KO male SCs post-injury compared to sham controls (Figure 4I-J). Quantification of Sox10+ cells revealed a significant decrease in *Ndufb5* expression in WT male SCs post-injury, while Sarm1KO cells showed no significant change (Figure S5D,E). Notably, mitochondrial respiration gene expression did not correlate with any specific injured cluster (Figures S4J, S5A-C), but were more highly expressed in Sarm1KO 1dpi SCs (Figure S5K). Additionally, Sarm1KO SCs in females showed a more pronounced upregulation of mitochondrial respiration genes at 1dpi than in males (Figure S6A, Supplementary Table 7), suggesting sex differences in the glial response to peripheral nerve injury. X-Y scatter plots of significantly regulated genes (adjusted p-value < 0.05) showed that most genes were regulated in the same direction in both males and females (purple, Figure S6E, F). However, WT SCs displayed some sex-divergent gene regulation (Figure S6E), with certain genes upregulated in one sex but downregulated in the other (orange). Notably, this sex-divergent pattern was not observed in Sarm1KO SCs (Figure S6F), where all significantly regulated genes showed concordant responses between sexes.

Interestingly, we did not observe induction of mitochondrial respiration genes at 2 hours post-injury in WT SCs (Figure S6B, Supplementary Table 8). Our pseudobulk analysis detected 485 differentially regulated genes at 2 hours post injury (Supplementary Table 8), with the top upregulated genes being *Ligp1*, *Ptx3*, and *Ctnna5* and downregulated *Cd44*, *Prx*, *Mbp.* These results suggest that SCs respond transcriptionally to injury quickly, but this does not depend on changes in mitochondrial respiration.

Overall, our results demonstrate that Sarm1 regulates SC metabolic responses to injury. Importantly, this metabolic regulation is independent of whether SCs progress along the transdifferentiation pathway (sc1, sc6) or remain in the less transdifferentiated state (sc4). The opposite regulation of mitochondrial respiration pathways in WT and Sarm1KO SCs highlights Sarm1’s role in shaping the SC injury response.

### Sarm1 regulates injury-induced metabolic shift away from mitochondrial respiration

Our snRNA-seq results suggested that WT and Sarm1KO SCs differ in their preferred metabolic pathway after injury. To investigate further, we performed mitochondrial bioenergetics assays using a Seahorse analyzer. Primary SCs underwent a mitochondrial stress test with sequential treatment with oligomycin, BAM15, antimycin A, and rotenone, to assess their mitochondrial energetic capacity under basal and stress conditions. To mimic the effects of injury on SCs *in vitro*, we activated ErbB2 signaling by adding Neuregulin 1 (Nrg1) for 24 hours, as previously described(*8*).

Consistent with previous observations(*8*), Nrg1 treatment did not significantly alter basal, maximal, or ATP-linked oxygen consumption rate (OCR) in WT SCs (Figure 4K, Figure S7A-C). In contrast, Sarm1KO SCs showed a significant increase in maximal OCR following Nrg1 treatment, while basal and ATP-linked OCR remained unchanged (Figure 4K, Figure S7A-C). This enhanced mitochondrial respiratory capacity suggests that in the absence of Sarm1, SCs maintain greater potential for mitochondrial respiration under stress conditions mimicking injury.

To validate that Nrg1 treatment recapitulates injury-induced transcriptional changes, we performed bulk RNA sequencing on cultured SCs. Nrg1-treated Sarm1KO SCs upregulated mitochondrial respiration genes in culture, mirroring the response observed after nerve injury *in vivo* (Figure S7D), while WT SCs showed minimal mitochondrial gene upregulation in either condition. Beyond these metabolic genes, broader genome-wide comparison revealed more modest concordance between *in vitro* and *in vivo* systems: 39-44% overlap in the top 1,000 differentially expressed genes, improving to 66-71% for the top 10,000 genes, with Sarm1KO showing consistently better agreement than WT (Figure S7E). These transcriptional findings corroborate the functional Seahorse data and validate Neuregulin treatment as an effective model for investigating metabolic aspects of the SC injury response, while highlighting that this *in vitro* system captures specific but not all features of the complex *in vivo* injury environment.

## Discussion

### Sarm1 Gates the Transition from Protective to Repair Schwann Cell States

Our findings reveal that Sarm1 functions as a molecular gate controlling SC state transitions following peripheral nerve injury. We propose that SCs adopt a transient Protection-Associated Schwann Cell (PASC) state characterized by intermediate pseudotime position, expression of Ptgds, Ptprd, Trf, and Fth1, and enrichment in Sarm1KO SCs at 1 day post-injury - immediately after injury, before progressing to the canonical Repair SC phenotype(*1*, *2*). This temporal gating function has critical implications: while early PASC activity supports axonal survival, timely progression to the Repair state is essential for successful long-term regeneration.

We propose a temporal model (Figure 5A) where Sarm1 operates with distinct kinetics across cellular compartments: glial Sarm1 contributes to early injury responses (hours), while neuronal Sarm1 acts continuously and more robustly throughout axon degeneration (days). Supporting this framework, glial-specific dSarm knockdown in *Drosophila* delays axon degeneration at 8 hours post-injury (Figure 1G,H), while neuronal Sarm1 knockdown produces stronger effects at 3 days (Figure S8A-B). This temporal separation explains why SC contributions are detectable in acute experimental windows but masked in longer-term experiments when neuronal effects predominate. Importantly, we recognize that the relative contribution of glial Sarm1 to axon degeneration is substantially smaller than the well-established neuronal Sarm1 phenotype(*13*), but becomes apparent when examining appropriate temporal windows and employing cell-type-specific manipulation.

**Figure 5:**
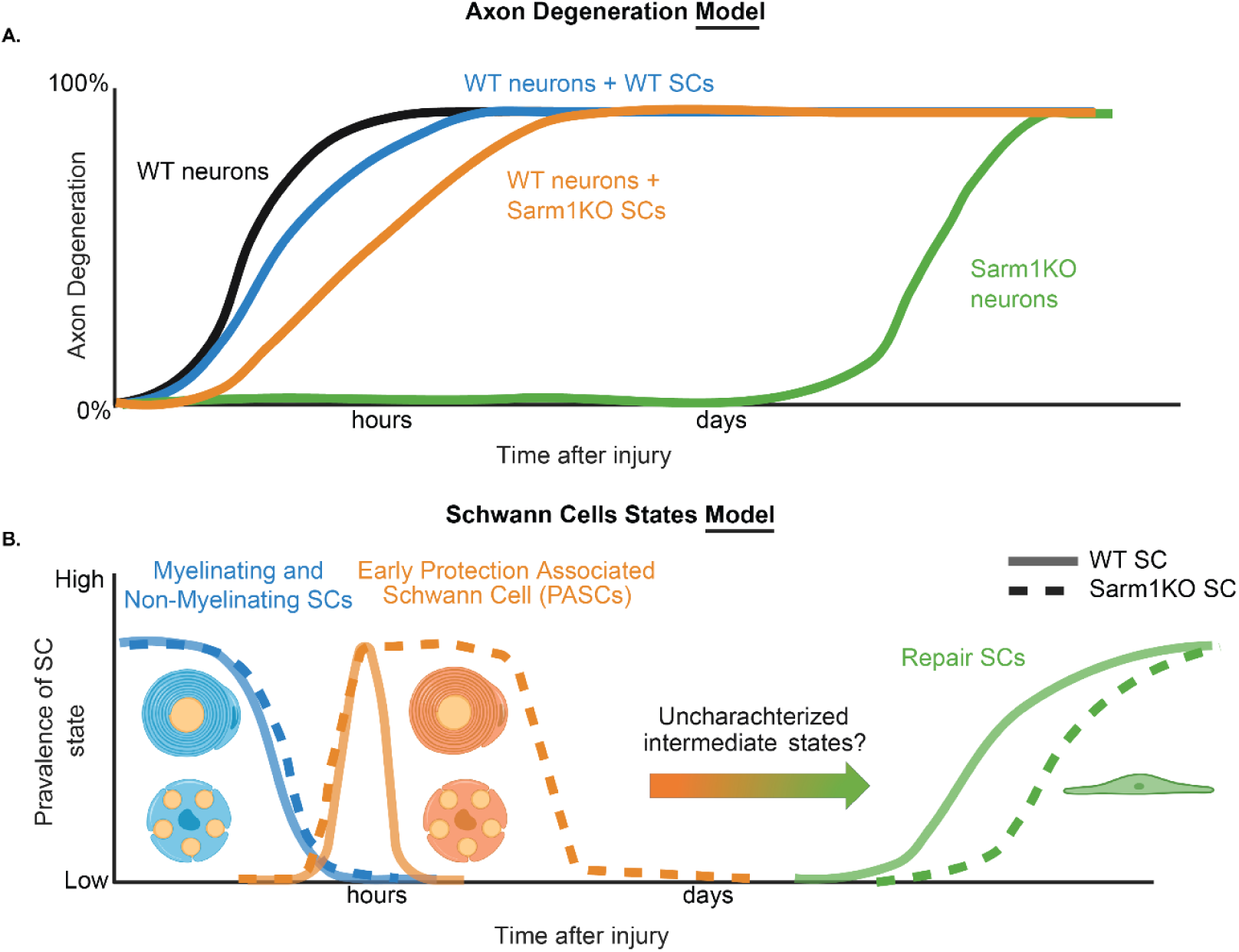
Axon degeneration and Schwann cell state dynamics following nerve injury in wildtype and Sarm1KO models. **A.** Axon Degeneration Model. Time course of axon degeneration following nerve injury in wildtype (WT) neurons alone (black), WT neurons co-cultured with WT Schwann cells (SCs; blue), WT neurons co-cultured with Sarm1KO SCs (orange), and Sarm1KO neurons alone (green). WT neurons undergo rapid degeneration within hours after injury, which is partially delayed by co-culture with WT SCs and further delayed by Sarm1KO SCs. Sarm1KO neurons show no significant degeneration for several days post-injury before eventually degenerating. **B.** Schwann Cell States Model. Working model of SC state transitions following nerve injury in WT (solid lines) and Sarm1KO (dashed lines) conditions. Following injury, myelinating and non-myelinating SCs (blue) rapidly downregulate their homeostatic programs. In WT conditions, early protection-associated Schwann cells (PASCs; orange) transiently emerge within hours before transitioning to repair SCs (green) over the course of days. We propose that in Sarm1KO conditions (dashed lines), the window of the PASC state is extended, potentially altering the dynamics and prevalence of subsequent SC states. The y-axis represents the relative prevalence of each SC state over time.

### SC-Derived Sarm1 Contributes Non-Cell-Autonomously to Axon Degeneration

A central finding of this work is that Sarm1 acts cell-autonomously within SCs to influence axon degeneration extrinsic to the axon itself. Using three independent experimental systems — an *in vitro* microfluidic co-culture, a mouse SC-conditional knockout, and a Drosophila pan-glial knockdown model — we consistently find that reducing SC or glial Sarm1 confers protection to axons after injury.

The field has historically studied Sarm1 in neuronal cultures and full knockout animals — contexts that either lack SCs or cannot distinguish cell-type-specific contributions. The sufficiency of axon-intrinsic Sarm1 loss to protect axons is well established, but sufficiency does not preclude contributions from the surrounding glial environment. Our results demonstrate that such a contribution exists and is conserved from flies to mammals. SC-derived Sarm1 acts as an early accelerant of degeneration rather than an independent executioner, most apparent in acute experimental windows before axon-intrinsic Sarm1 activity dominates.

This raises an important question: what state are SCs in when this early protective or degenerative activity occurs, and what molecular programs underlie it?

### PASC: A Transiently-Occupied Protective State

The transcriptomic data point to a previously uncharacterized SC state that emerges immediately following injury, preceding the canonical repair program. We term this the Protection-Associated Schwann Cell (PASC) state, defined by an intermediate pseudotime position between homeostatic and repair-like clusters, expression of genes distinct from both myelinating and repair programs — including Ptgds, Ptprd, Trf, and Fth1 — and a functional capacity to delay axon degeneration. Our single-nucleus RNA sequencing at multiple timepoints after injury, include 45,176 total nuclei with 7,082 injured SC nuclei, revealed that Sarm1KO SCs are enriched in cluster sc4 at 1 day post-injury (Figure 3C, Supplementary Table 2). Pseudotime analysis places sc4 as an intermediate state in both genotypes (Figure 3H-I).

The challenge in detecting PASC in wild-type animals likely reflects the rapid kinetics of this transition. While WT SCs transit through this state quickly—transiently appearing in cluster sc4 at 1 day post injury but enriched in the transcriptionally similar cluster sc2—many Sarm1KO SCs accumulate in sc4, suggesting a bottleneck at this transition point. Analysis at 2 hours post-injury revealed that WT SCs are already initiating transdifferentiation into cluster sc0, consistent with rapid transcriptional responses that precede the metabolic shifts observed at 24 hours. We propose that SCs may transiently adopt a PASC-like state during normal injury responses, though this transition occurs so rapidly in WT animals that it is difficult to capture without perturbation (Figure 5B). Sarm1 deletion slows this transition, allowing us to characterize this otherwise fleeting state. Importantly, substantial numbers of Sarm1KO SCs still progress to transdifferentiated states (sc1, sc6), indicating that Sarm1 deletion delays rather than completely blocks SC responses.

The existence of a PASC checkpoint likely reflects an adaptive strategy. The Repair SC phenotype involves transdifferentiation, active myelin breakdown, and debris clearance—processes that, while essential for regeneration after severe injury, could cause unnecessary damage to salvageable tissue. By maintaining a state that expresses protective factors while retaining differentiated characteristics (myelin-associated genes), SCs may provide early axonal support during the critical window when axon fate remains undetermined. Only once and if degeneration becomes inevitable does Sarm1-mediated checkpoint passage trigger the transition to Repair SCs. This model predicts that PASC maintenance duration should correlate inversely with injury severity, though testing this hypothesis it will be necessary to develop methods that allow for independent manipulation of injury severity or to utilize models of neurodegeneration, such as peripheral neuropathies, along with monitoring Sarm1 activity.

### PASC-Mediated Protection: Mechanisms and Metabolic Adaptations

The metabolic alterations in Sarm1KO SCs—enhanced oxidative phosphorylation and sustained expression of TCA cycle genes—represent a consistent phenotype across our experimental systems. Whether this metabolic profile directly contributes to the protective phenotype or reflects altered cellular homeostasis consequent to Sarm1 loss requires further investigation. Elevated NAD+ levels observed in Sarm1-deficient macrophages(*30*) and monocytes(*31*) raise the possibility that Sarm1 modulates NAD+ levels in SCs, influencing their metabolic state and function. Additionally, Sarm1’s known role in glial phagocytosis(*32*) suggests that changes in phagocytic receptor composition in PASCs could alter the balance between debris clearance and trophic support. Multiple molecular mechanisms likely operate simultaneously to mediate PASC-dependent protection, with their relative contributions and potential synergies remaining to be determined through targeted functional studies.

Another potential mechanism at play is through Fth1. Upregulation of transcription of this gene in cluster putative PASC cluster - sc4 represents one potential contributor. Fth1 catalyzes ferrous iron oxidation and sequestration, preventing iron-catalyzed reactive oxygen species generation(*28*). Recent work demonstrated that oligodendrocytes release Fth1 via extracellular vesicles, delivering protective cargo to neurons and conferring resistance against ferroptotic cell death(*29*). Whether SCs employ similar transcellular Fth1 transfer mechanisms, and whether this contributes to the axon protection we observe, requires direct experimental validation.

### Integration with Recent Findings

Our work captures the acute injury response (hours to 1 day) and complements recent studies examining later timepoints. Giger and colleagues demonstrated that Sarm1KO mice show reduced c-Jun and p75NTR induction, impaired SC transdifferentiation, and delayed myelin clearance at 3-7 days post-injury(*19*). Their nerve grafting experiments elegantly demonstrated that WT axons regenerate poorly into Sarm1KO grafts, while Sarm1KO axons regenerate well into WT grafts, confirming that the regeneration deficit stems from the distal nerve environment rather than intrinsic axonal properties. Thus, while we demonstrate that Sarm1 influences early protective responses (hours to 1 day), their work reveals consequences of disrupting the subsequent transition to the repair program (3-7 days). Prolonged PASC maintenance provides short-term protection but prevents the timely transition to Repair SCs necessary for myelin clearance and regeneration support (Figure 5).

The temporal integration of our findings with Babetto et al.’s work(*8*) reveals a more complete picture of SC metabolic dynamics. They demonstrated that WT SCs undergo a dramatic glycolytic shift at 24-36 hours post-injury driven by mTORC1 signaling, which enhances bioenergetic support to injured axons through increased lactate production. Our data showing that WT SCs did not upregulate mitochondrial respiration genes at 2 hours post-injury (Figure S6B) suggests these metabolic programs operate in distinct temporal windows. We propose a model where initial injury responses (0-2 hours) involve transcriptional changes without major metabolic reprogramming, followed by potential oxidative metabolism (captured in our PASC state at hours to 1 day), and subsequent glycolytic activation (24-36 hours, as shown by Babetto et al.(*8*)). The Sarm1-mediated transition may coordinate progression through these metabolic states.

The apparent contrast with Sundaram et al., who reported persistent increases in oxidative phosphorylation genes and metabolic activity in WT SCs from 1 week to several weeks after crush injury(*33*), likely reflects differences in injury models and temporal windows. In crush injuries where axon-glial interactions persist, prolonged oxidative metabolism may support both protection and eventual regeneration. In transection injuries, the rapid changes observed in Sarm1KO SCs may reflect different functional demands. This interpretation suggests that metabolic programming must be matched to injury context, with distinct optimal strategies for different injury severities and temporal phases.

### Broader Implications

Together, our findings establish Sarm1 as a multi-functional regulator of early SC injury responses. By combining single-cell transcriptomics with functional validation across species, we demonstrate that Sarm1 gates SC state transitions during the critical early hours post-injury. This gating function coordinates metabolic programming, putative protective factor expression, and the timing of repair program activation—revealing unexpected complexity in the SC response to nerve injury. By integrating our results with recent work demonstrating Sarm1’s cell-autonomous functions in macrophages and monocytes(*30*, *31*) and its requirement for timely SC repair program activation(*19*), a more complete picture emerges: Sarm1 operates as a multi-cellular coordinator of injury responses, with distinct temporal and functional roles in neurons, glia, and immune cells.

Our findings offer a potential intervention point for peripheral nerve injuries and neurodegenerative diseases. While Sarm1 inhibitors are being developed primarily to target neuronal Sarm1, our results demonstrate that Sarm1 also regulates critical processes in neighboring SCs. This raises considerations for therapeutic development: systemic Sarm1 inhibition will affect not only the intended neuronal targets but also SC state transitions, which could have both beneficial and deleterious consequences depending on timing and context. While acute Sarm1 inhibition may prolong the protective PASC state and extend the window for axon preservation, long-term deletion impairs regeneration(*19*). This suggests pulsed or reversible Sarm1 inhibition may be more effective than sustained blockade. Alternatively, combinatorial approaches that temporarily stabilize the PASC state while simultaneously promoting repair programs through orthogonal pathways (e.g., c-Jun activation(*6*)) might capture the benefits of both phases without the liability of prolonged checkpoint arrest. The identification of specific PASC markers, particularly those associated with protective functions also opens possibilities for therapeutic strategies that provide neuroprotection without requiring direct Sarm1 manipulation or disrupting normal SC state transitions.

### Study Limitations and Future Directions

Several limitations warrant acknowledgment. First, while our Drosophila pan-glial knockdown validates a non-neuronal contribution to axon degeneration, it does not resolve the relative contributions of specific glial subtypes. We address this limitation through a combinatorial approach, using SC-conditional Sarm1 knockout in mice alongside Sarm1KO SC-WT neuron co-cultures to establish SC-autonomous effects more directly. Second, our single-nucleus sequencing provides temporal snapshots rather than continuous trajectories of SC state transitions. Higher-resolution temporal profiling and lineage tracing will be necessary to fully map the PASC-to-Repair transition and determine whether all SCs transit through cluster sc4 or whether parallel trajectories exist – a possibility raised by the observation that only a subset of SCs express detectable Sarm1. Additionally, because our snRNA-seq studies used full Sarm1KO rather than SC-conditional knockout mice, we cannot entirely exclude the possibility that Sarm1 loss in other cell types indirectly influences SC transcriptional responses. Third, while we demonstrate Sarm1 mRNA expression in SCs by RNA FISH and RT-PCR, protein detection remains challenging due to low expression levels and current antibody limitations. For a catalytic enzyme like Sarm1, even modest expression can produce substantial downstream effects through signal amplification, and our functional data across multiple models support physiologically meaningful activity.

Looking forward, targeted manipulation of markers enriched in the putative PASC cluster sc4 could provide mechanistic insight into how SC-derived Sarm1 accelerates axon degeneration. Direct NAD+ measurements in WT versus Sarm1KO SCs will be important for determining whether NAD+-mediated metabolic regulation underlies the phenotypes we observe.

## STAR Methods

### EXPERIMENTAL MODEL AND STUDY PARTICIPANT DETAILS

#### Animals

Mice: All mice are on a C57BL/6J.129S mixed background. C57BL/6J (referred to as WT), Sarm1 knockout (referred to as Sarm1KO) and Sox10 Cre animals were purchased from Jackson Labs. All snRNA-seq experiments were carried out in adult (3-6 month old) mice. Sarm1 flox mice were a generous donation by Jing Yang lab from Peking University.

Drosophila Melanogaster: stocks and experimental crosses were maintained on standard molasses formulation food (Archon Scientific) at 25°C.

#### *In Vitro* Experiments

Superior Cervical Ganglia (SCG) Culture: Mouse pups aged P1-P2 were dissected to isolate superior cervical ganglia (SCG). SCGs were plated at a density of 1 SCG in each compartmentalized microfluidic device (MFD) with 5um grooves as described earlier (*34*). Neurons were cultured in DMEM supplemented with 10% FBS, penicillin/streptomycin (1 U/mL), and 45 ng/mL of NGF derived from mouse salivary glands. Contaminating glia were removed using 10 µM cytosine arabinofuranoside (Sigma-Aldrich, catalog 10931) for 48 hours. Cultures were maintained until days 5-7 and used only if they had successfully extended axon projections to the opposing side of the MFD.

##### SC Culture

Mouse pups aged P1-P3 were dissected to isolate sciatic nerves. SC isolation and culture were performed as described earlier(*35*) with some modifications. Sciatic nerves were isolated and enzymatically digested in 0.05% Type I Collagenase (Worthington Biochemical Corp, catalog LS004196) for 30 minutes, followed by 30 minutes in 0.125% Trypsin (Millipore T4799-5G). Nerves were further mechanically dissociated by trituration. After centrifugation at 300 xg for 5 minutes, cells were purified using a 70 µm filter (Corning, catalog 352350) and plated at about 30,000 cells/cm^2^. Cells were cultured in MEM supplemented with heat-inactivated Normal Horse Serum (Invitrogen, catalog 26050088) and 1:100 Pen-Strep (Gibco, catalog 15070-063), 2 µM Forskolin (Sigma, catalog F3917), and 10 µM Neuregulin (PeproTech, catalog 100-03). The next day, the media was changed and supplemented with 10 µM cytosine arabinofuranoside (Sigma-Aldrich, catalog 10931) for 24 hours, as described earlier, to help remove proliferating fibroblasts (*36*). After that, cytosine arabinofuranoside was removed, and cells were maintained for up to a week until ready for experiments.

##### SCG-SC Co-Culture

4-5 days after SCG plating, SCs were removed from the dish using 0.05% Trypsin-EDTA and plated into the distal axon side of SCG MFDs at 10,000 cells/MFD. Cells were allowed to establish connections with axons for 1 day. Then, all the cell body side of microfluidic devices was aspirated as described earlier (*21*). 8 hours post-injury, cells were fixed in 4% paraformaldehyde for 20 minutes, washed 3 times in PBS, blocked in 5% donkey serum, 0.05% Triton-X-100 in PBS for 1 hour at room temperature. This was followed by primary antibody staining overnight at 4°C (Tuj1 (1:1000, made on site) and Sox10 (1:150, ThermoFisher, catalog PA5-37890). The next day, cells were washed 3 times in PBS, followed by a 1-hour incubation in Donkey anti-Mouse Alexa Fluor 568 (Fisher, catalog A10037) and Donkey anti-goat Alexa Fluor 488, then washed 3 more times in PBS. A mounting medium with Dapi Fluoromount-G (SouthernBiotech, catalog OB010020) was applied. Distal axon compartments were imaged at a consistent distance from the injury site across the full height of the microfluidic devices. All images were captured immediately adjacent to the device grooves (Figure 4A) to maintain consistency between experimental conditions.

The imaging was done on a Zeiss LSM 780 confocal microscope in the W. M. Keck Center at the University of Virginia.

## METHOD DETAILS

### Sciatic Nerve Transection Surgeries

Mice were anesthetized with isoflurane at 4-5% and then maintained via continuous inhalation at 1-2%, and eye lubrication was provided. Fur was removed from the right side of the back/hindlimb region, and the area was cleaned with 3 alternating wipes of iodine and 70% alcohol. Prior to incision, bupivacaine was applied locally (0.001 mL/gram mouse body weight, 0.25% solution) to prevent pain at the surgery site. An incision was made through the skin over the dorsal hindlimb region. Blunt dissection through the muscle allowed access to the sciatic nerve, which was transected with a pair of sterile surgical scissors, and a 2mm nerve segment was removed. The gluteal muscles were then brought back into their original anatomical position and secured in place with surgical sutures, and the skin was reapproximated by surgical sutures. The animal was also administered ketoprofen for pain management (0.004 mL/gram mouse body weight, 1 mg/mL concentration).

### Tissue Harvest

Tissue was collected at multiple timepoints: WT mice at 2 hours post-injury, WT and Sarm1KO mice at 1 day post-injury, and WT mice at 2 days post-injury. After surgery, mice were rapidly cervically dislocated without anesthesia for tissue collection to avoid confounding factors of anesthesia or CO_2_ on the rapidly changing transcriptome of the already injured sciatic nerve. Uninjured (sham) and injured sciatic nerves were immediately collected, avoiding a 2 mm stump at the injury site, and uninjured or sham nerves (left side) were collected. Visual confirmation that surgery yielded transection (rather than crush) and that the ends did not rejoin was also performed prior to harvest. Distal to cell body sciatic nerves were collected, placed in a vial, and immediately flash-frozen in liquid nitrogen. Tissue was stored at -80 °C until further processing.

### Nuclei Isolation, Library Preparation, and Sequencing

Tissue was defrosted, and 2-6 nerves per condition were pooled. Twelve experimental conditions were analyzed (detailed in Supplementary Table 9): WT and Sarm1KO (male and female) at sham and 1dpi, plus WT (male and female) at sham and 2hpi For the 1 day post injury experiments (Batches 1-4), sciatic nerves from multiple animals (2-6 animals) of the same sex were pooled to obtain sufficient material for each sample. Each batch consisted of n=2 samples (one pooled male sample and one pooled female sample), with both sham and 1dpi conditions from each genotype (WT and Sarm1KO) processed simultaneously. The experiment was repeated across 4 independent batches For the 2 hour post injury experiment we included 4 biological replicates (in “Batch 7”) - 2 male and 2 female (Table 9). Batch 5 contained only nuclei from Naive samples - sciatic nerves came from animals that did not go through injury (only used for clustering of all cells, not differential gene expression analysis). Pooled nerves from each condition were dounce homogenized, and nuclei were purified by density gradient centrifugation as described earlier (*37*, *38*) and re-suspended in buffer (molecular-grade PBS with 1% BSA, 0.1% Protector RNase inhibitor, 2mM MgCl_2_). Following that, Total Seq anti-Nuclear Pore Complex Proteins Hashtags (BioLegend, catalog numbers: 682209, 682211, 682213, 682205, 682207, 682215, 682219) were applied to track the sample-of-origin for each cell nucleus. Samples were then stained with NucBlue (Invitrogen, catalog R37605), and batches 4 and 5 were also stained with Donkey anti-Mouse IgG (H+L) Highly Cross-Adsorbed Secondary Antibody, Alexa Fluor 647 (ThermoFisher, catalog A-31571, 1:100) that binds to Total Seq antibodies to increase the purity of the nuclear fraction. The first batch was sorted on a Sony SH800 Cell Sorter, while the rest were sorted based on NucBlue+/AlexaFluor647+ nuclei on a Becton Dickinson Influx cell sorter. To ensure a clear fraction, we first gated based on a highly fluorescent Alexa Fluor 647 signal (Figure S2A), followed by selecting cells of appropriate size and complexity using forward vs. side scatter distribution. We then ensured that only singlets were selected by plotting trigger pulse width vs. forward scatter. Lastly, we selected a fraction with high fluorescence of the DNA intercalator. With these settings, nuclei were sorted in 18.8 µL of RT Reagent B from the 10x Genomics Chromium Next GEM Single Cell 3’ Kit v3.1. The remaining components of the Step 1 master mix reagents were combined in the capture tube, along with sufficient resuspension buffer to achieve a total volume of 75 µL. Subsequently, we followed the manufacturer’s instructions (10X Genomics, CG000204 Rev D) to process the sample into complementary DNA (cDNA) sequencing libraries. For the hashtag oligonucleotide (HTO) libraries, we followed a previously published protocol (*39*), with a modification: double-sided SPRI (solid-phase reversible immobilization) was performed twice to effectively separate HTO libraries from cDNA libraries after amplification and, if necessary, just before pooling libraries for sequencing. The first 3 batch libraries were then sequenced on the Illumina NextSeq 550 using a 75-cycle, high-output kit, while the last 2 batches were sequenced with NextSeq 2000 using a P2 100-cycle run following the manufacturer’s instructions.

### Data Analysis of Sequencing Data

We used the 10X Genomics Cell Ranger pipeline (version 9.0.1) to align reads to the mouse reference transcriptome (mm10-2020-A) and calculate gene-level expression values with Unique Molecular Identifier (UMI) correction. The inclusion of introns was enabled using the Cell Ranger include-introns parameter. We set the CellRanger force-cell parameter to 15,000 cells. To ensure only cells containing nuclear RNA were included in the downstream analysis, we employed hashtag oligonucleotide (HTO) demultiplexing using the MULTIseqDemux function. This hashtagging approach also enabled multiplexing of samples within a single 10x Genomics chip lane.

We incorporated TotalSeq indexes into sham sample hashtag libraries to distinguish between sham and injured cell nuclei. During demultiplexing, reads that did not align to sham hashtags were assigned to the injured condition based on their corresponding cell barcodes. To prevent misassignment due to potential barcode overlap between 10x Genomics lanes, we excluded cells with identical barcodes appearing across multiple lanes from the injured samples (detailed procedures available in the accompanying code). Additionally, we implemented quality control filters for each library, removing cells that fell more than 2 standard deviations below the mean for two key parameters: number of genes detected per cell (nFeature_RNA) and total number of molecules detected per cell (nCount_RNA). Cells containing more than 5% of reads from mitochondrial genes were excluded from the downstream analysis. Libraries with HTOs underwent filtering, where cells that expressed multiple HTOs (muliplets) or no HTOs (no binding occurred or ambient RNA) were eliminated from the analysis. Subsequently, all batches were integrated to correct for technical variance(*40*). We then log-normalized the data, selected the top 2,000 highest variance genes, integrated the libraries in Seurat using CCA integration method, scaled the data, conducted Principal Component Analysis (PCA) to reduce the dimensionality of the high-variance gene space, and clustered the cells using the Louvain algorithm based on Euclidean distance in the PCA space. For further subclustering just SCs, we did not integrate the data. Non-linear dimensionality reduction was performed through Uniform Manifold Approximation and Projection (*41*) for visualizing the clustered data in two dimensions. To select the most variable genes between genotypes and conditions, we employed the Edge Libra Differential Expression in R, which averages cells by batch within each cluster into a pseudo-bulk sample before performing differential expression analysis (*42*). To visualize cells as a function of pseudotime, we have utilized the Monocle3 (*43*) package and set sham cells as time 0. To detect genes enriched in each cluster we used Seurat’s FindAllMarkers (min.pct=0.25, logfc.threshold=0.25) function.

### RNA Fluorescence In Situ Hybridization (RNA FISH) Procedure

RNA FISH experiments were performed on sciatic nerves obtained as described in the “Tissue Harvest” section. However, the tissue was fresh frozen in Tissue-Tek® O.C.T. Compound (Sakura, catalog 4583) submerged in liquid nitrogen, cryo-sectioned on a cryostat (Thermo Scientific, Microm HM 525) at 10 µm thickness, and stored at -80 °C until ready to use. On the day before RNA FISH, slides were brought to room temperature and immediately incubated in 10% formalin for 20 minutes. They were then washed 3 times in PBS and left to dry overnight. The next day, sections underwent incubation for 10 minutes in hydrogen peroxide at room temperature, followed by Protease IV in a HybEZ II Oven at 40 °C. The tissue was then exposed to a 2 hour incubation with the probe of interest (Ndufb5, Ndufb10, Sox10, or Sarm1) at 40 °C. The tissue further underwent treatments with AMP1, 2, and 3, followed by HRP-C1, C2, C3, and HRP Blocker, components of (ACD Bio catalogs 323110, 322381). The fluorophores used to visualize probes of interest were TSA Plus FITC, Cy3, and Cy5 (Akoya Biosciences, catalog numbers NEL741001KT, NEL744001KT, NEL745001KT). Slides with tissue were covered with mounting medium DAPI Fluoromount-G (Fisher/SouthernBiotech, catalog OB010020) and coverslipped. Z-stack images were captured using a Leica Stellaris 5, and Zeiss 980 with a Z interval of 0.5 µm and the number of Z stacks being determined empirically for each image.

### Image Analysis

#### RNA FISH

To analyze the images, the Z-stacks were compressed using maximum intensity projection into a single plane image and imported into QuPath software. A mask over DAPI-positive nuclei was automatically selected to measure expression of transcripts in nuclei. To measure total cell expression, a 5 µm expansion around the nucleus was drawn automatically by the software to predict cytoplasm boundaries. The software was trained by a set of images selected by a blinded investigator to identify nuclei and cells positive for Sox10 probe (ACD, catalog 435931-C3) signal. The software was further trained to identify other probes (Ndufb5 (ACD, catalog 1303021-C1), Ndufb10 (ACD, catalog 1303041-C2), Ctnna2 (ACD, catalog 1303031-C2), and Sarm1 (ACD, catalog 504191-C2)). Eventually, the percentage of Sox10-positive cells also positive for probes of interest was quantified and compared between conditions using mean +/- standard error of the mean (SEM) and analyzed using unpaired t-test.

#### Immunocytochemistry

A blinded investigator quantified axon degeneration from fluorescence microscope images of cultured, Tuj1-immunostained axons as described earlier (*16*, *21*, *44*). In brief, randomly assigned 50 µm lines tracing along the length of axons, avoiding bundled axons. The investigator then counted the number of boxes containing 3 or more beads/blebs, categorizing them as degenerating axons. The percentage of total degenerating axons was calculated using Microsoft Excel. Data were represented as mean +/- standard error of the mean (SEM), two-way ANOVA with Tukey multiple comparisons test was performed to analyze differences between conditions. Each experiment was conducted a minimum of 3 times using separate litters of mouse pups as a separate replicate.

### Mitochondrial Stress Test

SCs were isolated from WT and Sarm1KO P0-P1 sciatic nerves and plated at 20,000 cells/well of an XFe96 cell culture microplate for 5-7 days as described above, with one difference: SCs were supplemented with 20 µg/mL of Bovine Pituitary Extract (Fisher Scientific, catalog 13-028-014) instead of Nrg. 24 hours before the assay, a set of wells was subjected to Nrg treatment (200 ng/ml), simulating injury. After Nrg treatment, the media was changed to mitochondrial stress test medium (Corning catalog 50-003-PB). Mitochondrial activity was assessed by measurement of O_2_ consumption rate on a Seahorse XFe96 instrument (Agilent Technologies). The rate of O_2_ change was measured every 13 minutes for a 3-minute interval before sequential challenge with 1) 1 μM oligomycin (Sigma Aldrich catalog 75351), 2) 2 μM BAM15 (Cayman Chemical Company catalog 17811), and 3) 10 μM antimycin A (Sigma-Aldrich catalog A8674) and 10 μM rotenone (Sigma-Aldrich catalog R8875-1G). Maximal OCR was measured as post-BAM15 OCR minus post-antimycin A/rotenone OCR. Basal OCR was measured as pre-oligomycin OCR minus post-antimycin A/rotenone OCR. Upon completion of the assay, cells were immediately fixed in 4% PFA for 20 minutes, washed 3x in PBS for 5 minutes and incubated in a DAPI mounting medium. Cells were imaged on a Zeiss LSM 780 confocal microscope in the W. M. Keck Center at the University of Virginia, cell quantification was performed using QuPath software. The values from mitochondrial activity measurements were normalized to cell numbers in each well. Data is represented as mean +/- standard error of the mean (SEM) with each data point representative of independent litters harvested, unpaired two-tailed t-tests were used to analyze differences between conditions.

### Bulk RNA sequencing

#### RNA collection

SCs coming from WT and Sarm1KO backgrounds were isolated from three independent litters and cultured as described above in 35mm dishes for 7 days. Once established, cultures were stimulated either with neuregulin (Nrg, 200 ng/ml) or left untreated for 24 hours using the same conditions as for the mitochondrial stress test.

At 24 hours post-treatment, mRNA was collected using the following protocol: 500 μl of TRIzol Reagent (Invitrogen, 15-596-026) was applied to each dish, and cells were extensively mixed with the reagent by pipetting. The suspension was transferred to a 1.5 ml microcentrifuge tube and incubated for 5 minutes at room temperature. Next, 100 μl of chloroform (Fisher Scientific, AC158210010) was added, and the mixture was vigorously shaken for 15 seconds followed by a 2-3 minute incubation at room temperature. Samples were then centrifuged for 15 minutes at 12,000×g at 4°C, and the upper aqueous layer was collected. For RNA precipitation, 250 μl of isopropanol was added to the aqueous layer and incubated for 10 minutes. Samples were centrifuged again for 10 minutes at 12,000×g at 4°C. The resulting white pellet was resuspended in fresh isopropanol and incubated overnight at 4°C

#### cDNA preparation and sequencing

cDNA library preparation was performed strictly according to manufacturer protocols. RNA was purified using the NEBNext Poly(A) mRNA Magnetic Isolation Module (New England BioLabs, E7490S). cDNA libraries were prepared using the NEBNext Ultra II RNA Library Prep Kit for Illumina (New England BioLabs, E7770S), and sequencing indexes were applied using NEBNext Multiplex Oligos for Illumina (Index Primers Set 1, E7335S).RNA purification was performed using SPRI Select beads (Beckman Coulter, B23317). RNA quality was assessed using the TapeStation High Sensitivity RNA ScreenTape (Agilent, 5067-5579). RNA Integrity Number (RIN) scores ranged between 7.1 and 10. The cDNA libraries were sequenced by Azenta using 2×150 bp paired-end reads, generating approximately 350M paired-end reads (∼30M reads per sample).

#### Analysis

The FASTQ files provided by Azenta were aligned to the mouse transcriptome “Mus_musculus.GRCm39.cdna” using Salmon 1.2.0 for indexing and quantification. Quantified data was imported into R using the tximport package. Differential expression analysis was performed using DESeq2, comparing untreated versus neuregulin-treated cells from WT versus Sarm1KO backgrounds. Refer to the accompanying code for detailed analysis parameters.

### RT PCR

#### Reverse transcription of Sarm1

RNA from bulk RNA sequencing preparation was reverse transcribed using the Luna Script RT SuperMix Kit (BioLabs, E3010S) according to the manufacturer’s instructions. Equal amounts of cDNA (∼100 ng) and diluted cDNA (∼50ng) were amplified using the REDExtract-N-Amp™ PCR ReadyMix Kit (SigmaAldrich, R4775). Each 10 μL reaction contained 5 μL REDExtract-N-Amp PCR ReadyMix, 1 μL cDNA template, 3.5 μL nuclease-free water, and 0.5 μL primer mix.

Primer for Sarm1 RNA were used as already described (Sarm1 forward GGACCATGACTGCAAGGACTG, reverse CCGTTGAAGGTGAGTACAGCC) and GAPDH (FisherScientific, 43-529-32E) was used as a loading control. Cycling parameters for both primers: 1. 95°C, 2:00 2. 95°C, 0:30 3. 60°C, 0:30 (Tm is 64.5) 3. 72°C, 0:30 4. Goto 2, 31X 5. 72°C, 10:00 6. 4°C, forever. PCR products were resolved on 2.7% agarose gels in 1× TAE buffer at 80V for approximately 1 hour and visualized using a ChemiDoc BioRad imager.

#### Analysis

Tight boxes were drawn across the bands coming from the both GAPDH and Sarm1 bands coming from more and less concentrated samples. The mean integrated density was calculated in Fiji ImageJ. The value of Sarm1 was divided by its corresponding GAPDH measurement to normalize the values based on the amount of RNA loaded. The values from both concentrations were averaged. The analysis was done in GraphPad prism.

### Transmission Electron Microscopy Conditional Knockout injury experiments

#### Tissue processing and imaging

8–12 week old mice of the following genotypes — Sox10 Cre/+; Sarm1fl/+ and Sox10 Cre/+; Sarm1fl/fl littermates — underwent transcardial perfusion with 4% paraformaldehyde and 2.5% glutaraldehyde 48 hours following sciatic nerve transection injury. Distal sciatic nerves were isolated and post-fixed in the same fixative solution. Tissue was processed for transmission electron microscopy by the Molecular Electron Microscopy Core (MEMC) at the University of Virginia. To avoid artifacts associated with the injury site, the first 1.5 mm of the injured nerve stump was excluded, and ultrathin 70 nm sections were collected by ultramicrotomy at 1.5 mm distal from the injury site. At least 10 fields of view per animal were imaged on an FEI Tecnai F20 at 120 kV.

#### Image analysis

A blinded investigator counted the total number of large caliber axons per field of view, excluding any axons not fully contained within the frame. Each axon was then assessed for signs of degeneration. The number of degenerating axons was summed across fields of view for each animal, with ∼ 100 -300 axons counted per animal, and the percentage of degenerating axons was calculated. Both sexes were included in the analysis, with equal numbers of male and female samples represented for each genotype.

To assess axon and myelin morphology, blinded investigators manually traced the outer and inner borders of the myelin sheath. Axon and axon and myelin diameter values was derived from the area to account for axons that are not perfectly circular. The g-ratio was calculated by dividing the inner diameter by the outer diameter of the myelin sheath.

### Drosophila Melanogaster wing injury experiments

#### Stocks

*Drosophila melanogaster* stocks and experimental crosses were maintained on standard molasses formulation food (Archon Scientific) at 25°C. On the day of eclosion, experimental flies were transferred to 29°C to increase RNA interference (RNAi) and maintained at this temperature for 6 days prior to injury experiments.

The following lines were used: RepoGal4 (*45*); UAS-dSARM RNAi (Vienna Drosophila Resource Center (VDRC) 102044); UAS-dSARM RNAi (VDRC 105369); UAS-Luciferase RNAi (Bloomington Drosophila Stock Center (BDSC) 31306, TRiP.JF01355); Dpr1 Gal4 (*46*); tubQS,hsFLP, FRT19A (BDSC 30130), w,hsFLP,FRT19A;Vglut-QF2,QUAS-CD8GFP,aseFLP2d/CyO;repo-Gal4,tub-Gal80ts,as eFLP3b/TM3[Sb,e] (*47*). See Table 10 for additional details on experimental crosses.

#### Wing Injury Assay

Flies were anesthetized with CO2 and subjected to wing transection injury in proximity to the LV0 vein on the LV1 vein, ablating all visible cell bodies as depicted in the schematic diagram (Figure 4D). Injuries were performed using sharp MicroPoint Scissors (EMS, VANNAS Scissors; angled on side, delicate, 5-mm cutting edge, #72933–04). The contralateral wing was left uninjured and served as a control.

At specified time points post-injury—8 hours for RepoGal4 (9-10 animals per condition) experiments and 3 days for Dpr1 Gal4 experiments (8-11 animals per condition)—both wings were collected and mounted on slides with Halocarbon Oil 27, covered with coverslips, and immediately imaged. Images were acquired using an Innovative Imaging Innovations (3i) spinning-disc confocal microscope equipped with a Yokogawa CSX-X1 scan head and 100× objective. Z-stacks were collected at 0.27 μm intervals through the entire thickness of the L1 vein, following established protocol (*47*) and shown in Figure 4D.

#### Quantifications

For the Dpr1 Gal4 experiments: a blinded investigator scored axon degeneration on a scale from 0 (least degenerated) to 6 (most degenerated) based on membrane signal integrity and the presence of axonal swellings.

For the RepoGal4 MARCM experiments: a blinded investigator quantified axonal breaks and puncta/swellings by counting these features at three different 30 μm segments along individual axons. When possible, at least two different axons were analyzed per image. In cases where only one axon was labeled or axons were too bundled for individual analysis, only one axon was counted. When multiple axons were quantified from a single image, the counts from each axon were averaged.

## QUANTIFICATION AND STATISTICAL ANALYSIS

Quantifications and statistical analysis were performed for different assays as described in methods and legends.

## Lead Contact

Further information and requests for resources and reagents should be directed to and will be fulfilled by the lead contact, Christopher Deppmann, Ph.D.

## Author’s contributions

E.S., J.C., S.H.-C., S.K.,N.L., J.C.-B.and C.D. planned all the experiments. E.S., C.C., J.L, C.P., A.T., T.V., J.G., ,C.K.-A.and S.H.-C. performed all the experiments and analyzed the data. E.S., C.D., J.C. wrote the manuscript with contributions from co-authors

## Acknowledgements

We thank members of the Deppmann, Campbell, Kucenas and Coutinho-Budd labs for helpful discussions. We are grateful to Bogdan Bierowski and Elisabetta Babetto for their insightful conversations that contributed to this work. We acknowledge the Keck Center for Cellular Imaging for the use of the Zeiss 780 and 880/980 microscopy systems [PI: AP; NIH-OD025156 and PI: AP; NIH-OD016446]. During the preparation of this work, the authors used Claude (Anthropic) for proofreading, grammar checking, and manuscript polishing, including line editing, consistency auditing, and formatting review. After using this tool, the authors reviewed and edited all content and took full responsibility for the content of the publication.

## Ethics approval

All animal care and experiments were performed in compliance with the Association for Assessment of Laboratory Animal Care policies and approved by the University of Virginia Animal Care and Use Committee.

## Data Availability

The RNA sequencing data and snRNA sequencing data generated in this study will be deposited in the GEO database under accession code [placeholder]. Other data are available from the corresponding author upon reasonable request. The code generated for analysis of this data can be found on Github repository https://github.com/stepanovacz/Sarm1_SchwannCell

---

**Supplemental Figure 1:**
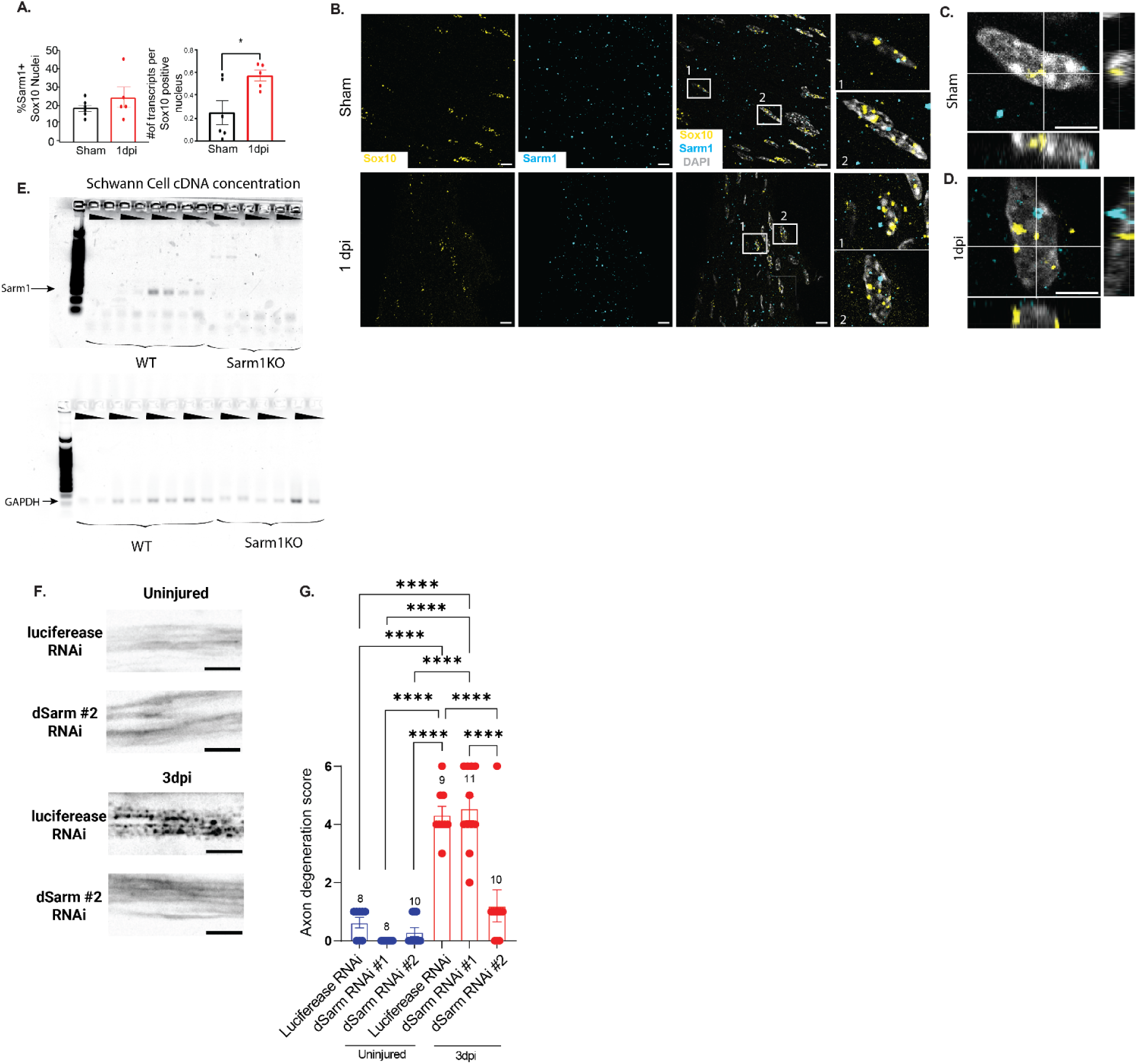
Sarm1/dSarm role in axon degeneration in SCs. **A.** Left: Quantification of the percentage of Sox10+ nuclei that are also Sarm1+. Right: Quantification of the average number of Sarm1 transcripts per Sox10+ nucleus. Data presented as mean ± SEM. Statistical comparison by unpaired two-tailed t-test, p-value * = 0.0262. **B.** Representative images of RNA in situ hybridization for Sox10, Sarm1, and merged with DAPI in sham sciatic nerves. Scale bar = 10 µm. Numbered boxes indicate regions shown in higher magnification insets. **C-D.** 3D reconstructions of SC nuclei showing the spatial relationship between Sox10 (yellow) and Sarm1 (cyan) transcripts in sham (C) and 1 day post-injury (D) sciatic nerves. Bottom: perspective view. Right: cross-sectional view. Scale bar = 5 µm. **E.** Full gel image showing all biological replicates for Sarm1 and GAPDH from RT-PCR (corresponding to cropped regions shown in Figure 1A). Triangles indicate cDNA concentration loaded (1× vs 0.5×). **F,** Representative images of CD8::GFP-labeled neurons with UAS-Luciferase and UAS-dSarm RNAi crossed to Dpr1-Gal4 driver. Scale bar = 10 µm. **G.** Blinded investigator-assigned degeneration scores ranging from 0-6 (0 being the least degenerated, 6 being the most degenerated). Number of biological replicates shown at the top of each bar graph. Data presented as mean ± SEM. Statistical comparison by two-way ANOVA with Tukey’s test for multiple comparisons, p-value **** < 0.0001.

**Supplemental Figure 2:**
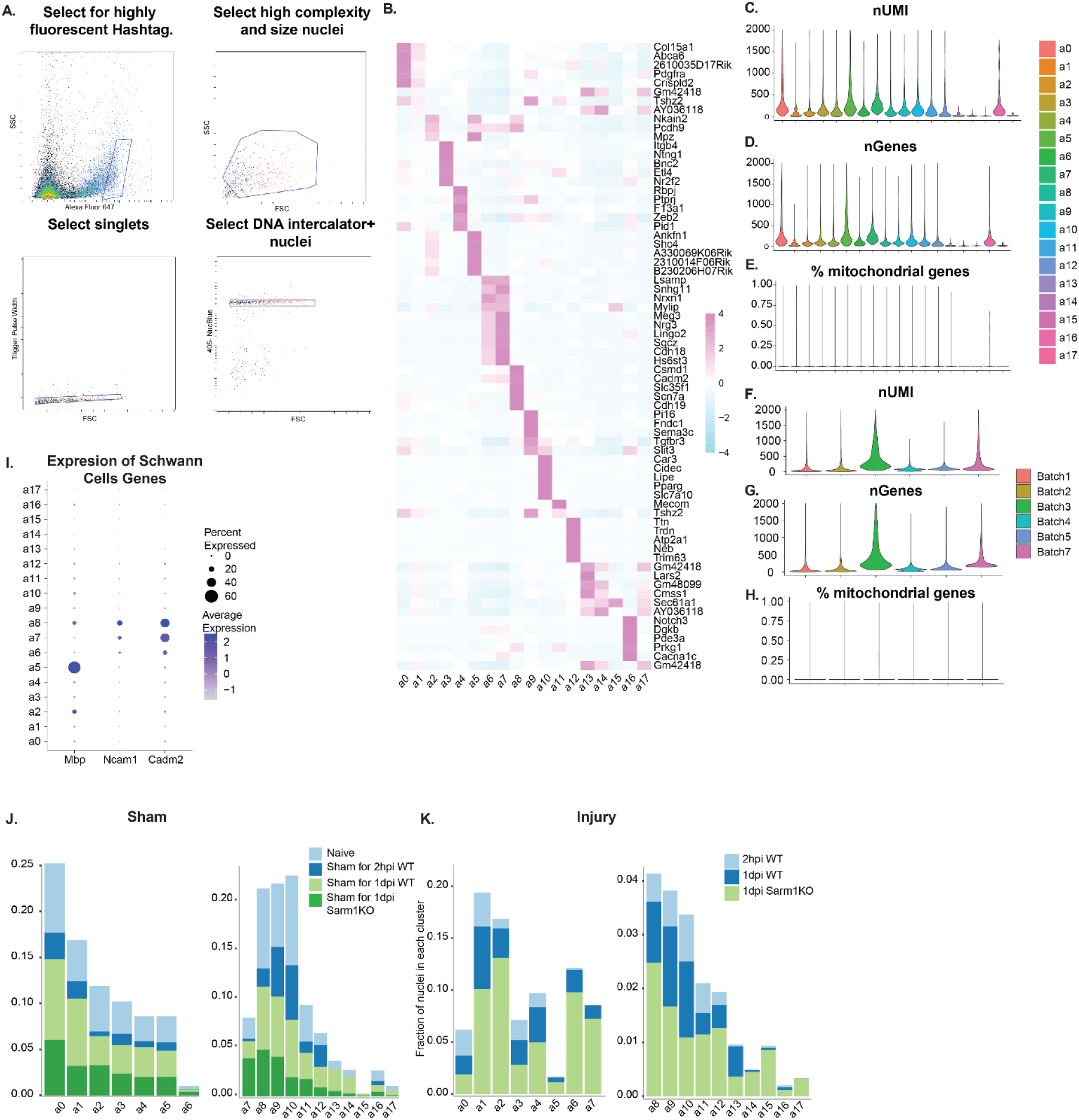
Supplemental Information on Single-Nucleus RNA Sequencing Analysis. **A.** Fluorescence-activated cell sorting (FACS) workflow for nuclei selection: 1)Selection of nuclei with high fluorescence for nuclear pore complex antibodies; 2) Gating for high complexity (SSC) and size (FSC) nuclei; 3) Selection of singlets; 4) Gating for DNA intercalator-positive nuclei. **B.** Heatmap showing cluster-level expression of top 5 marker genes for each identified cluster, based on relative expression levels. **C-H.** Cell quality metrics between clusters and batches. Distribution of UMIs (C and F), number of genes (D and G), and percentage of mitochondrial genes (E and H) per cluster (C-E) and per batch (F-H). **I.** Dot plot showing expression of Mbp, Ncam1, and Cadm2 on a per-cluster basis across all sequenced cells. **J-K.** Stacked bar plots showing the proportion of nuclei from each cluster in sham (J) and injured (K) conditions, split by experimental group (WT, Sarm1KO).

**Supplemental Figure 3:**
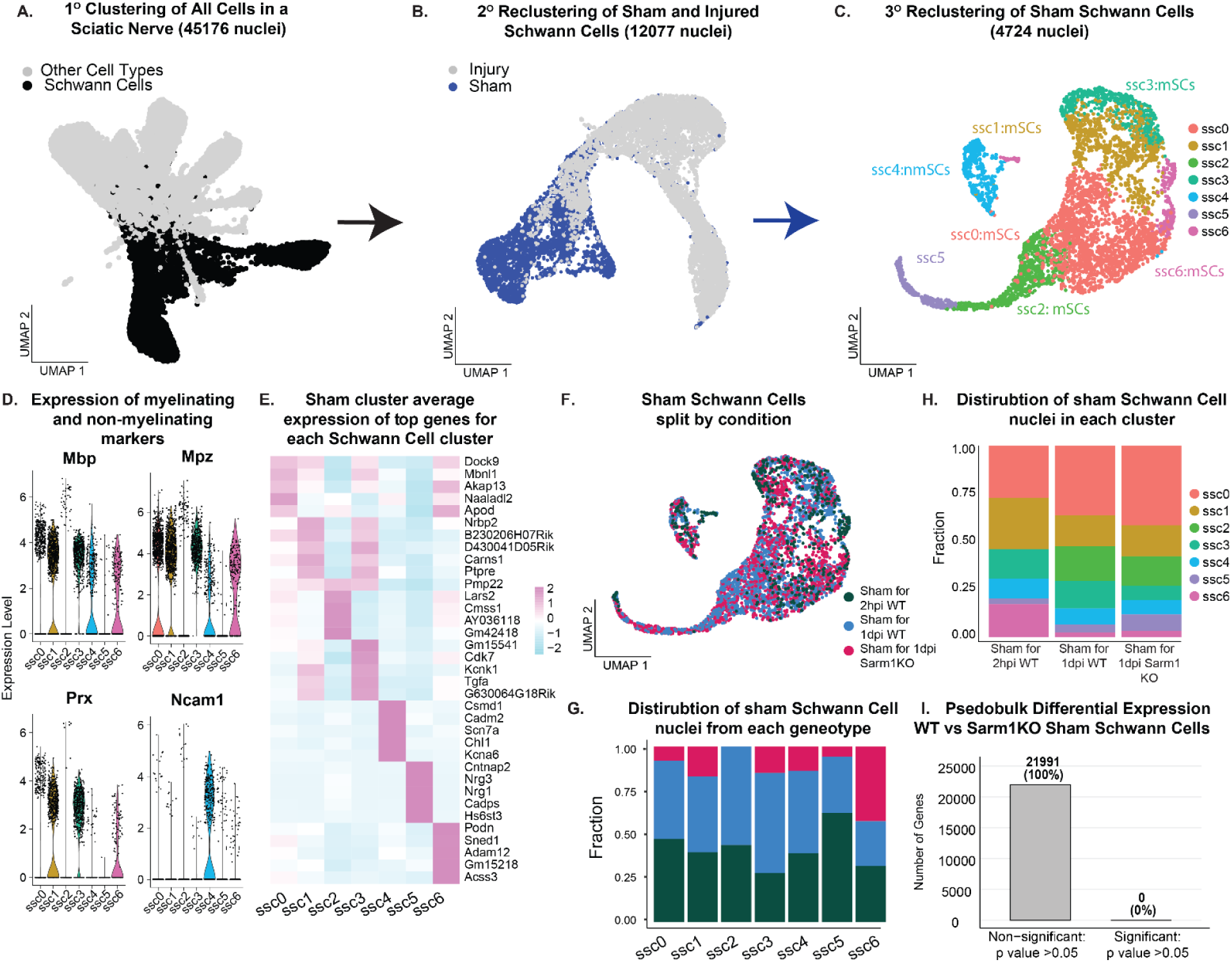
SCs in uninjured sciatic nerves of WT and Sarm1KO mice exhibit minimal differences in gene expression. **A.** UMAP showing selection of SC clusters a2,a5,a6,a7 and a8 (highlighted in black) as revealed by expression of Mbp, Ncam1 and Cadm2 for reclustering. **B.** UMAP of secondary clustering analysis performed exclusively on SCs from sham conditions (Sham SCs; blue). **C.** UMAP of tertiary clustering analysis on SCs from uninjured nerves, revealing 7 distinct clusters. **D.** Violin plot of myelinating and non-myelinating markers showing myelinating and non-myelinating SCs. **E.** Heatmap depicting average expression of top 5 genes significantly enriched in each cluster (adjusted p-value < 0.05). **F.** UMAP of Sham SCs color-coded by genotype (WT or Sarm1KO). **G.** Bar graph showing the proportion of WT and Sarm1KO Sham SCs in each cluster. **H.** Bar graph illustrating the distribution of clusters within WT (split by time to collection: 2 hours or 1 day after contralateral nerve surgery) and Sarm1KO Sham SC populations. **I.** Bar plot showing number and percentage of significantly (p < 0.05) and non-significantly (p > 0.05) regulated genes between WT and Sarm1KO SCs.

**Supplemental Figure 4:**
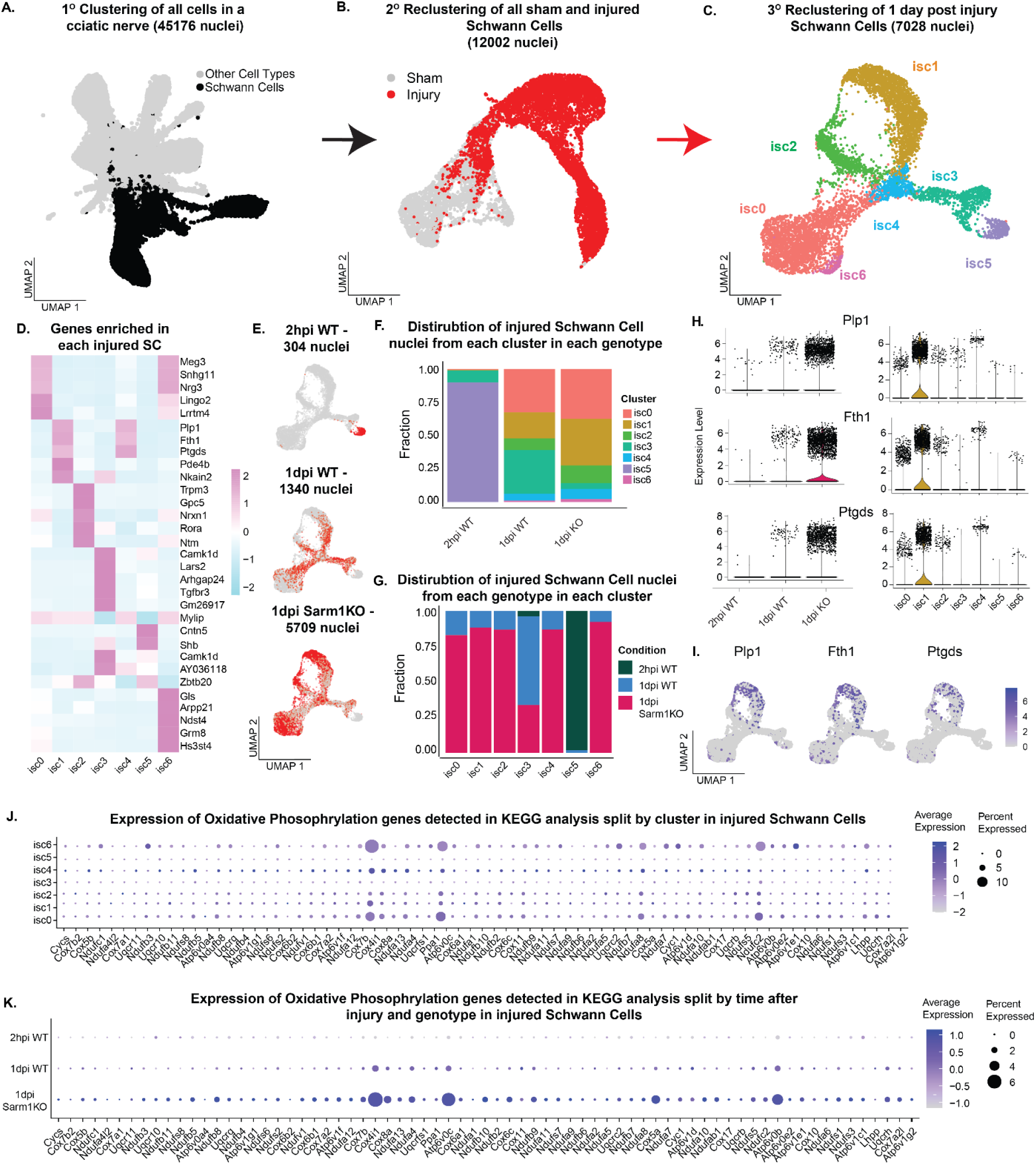
Supplemental analysis of injured SCs in WT and Sarm1KO mice. **A.** UMAP showing selection of SC clusters a2,a5,a6,a7 and a8 (highlighted in black) as revealed by expression of Mbp, Ncam1 and Cadm2 for reclustering. **B**. UMAP plot showing distribution of injured SCs. Red cells indicate the selected SCs. **C.** UMAP of tertiary clustering analysis on SCs from injured nerves, revealing 7 distinct clusters. **D.** Heatmap depicting average expression of top 5 genes significantly enriched in each cluster (adjusted p-value < 0.05). **E.** UMAP plots showing distribution of injured SCs, split by genotype (WT or Sarm1KO) and time after injury (2 hours post-injury (2hpi) and 1 day post-injury (1dpi)). Red cells indicate the selected SCs for the indicated genotype and timepoint, while gray indicates all injured SCs. Total nuclei counts are indicated for each condition. **F.** Stacked bar plot illustrating the proportion of injured SCs in each cluster, stratified by genotype and time after injury. **G.** Bar plot depicting the proportion of injured SCs from each experimental condition (WT and Sarm1KO, time after injury) within each cluster. **H-I.** Violin plots (H) and feature plots (I) showing expression of 3 different markers enriched in Sarm1KO SCs 1 day post-injury, showing particular enrichment in cluster is1. **J-K.** Dot plots showing expression of mitochondrial respiration genes (identified in Figure 3G) split by cluster (J) and by genotype and condition (2hpi, 1dpi) (K).

**Supplemental Figure 5:**
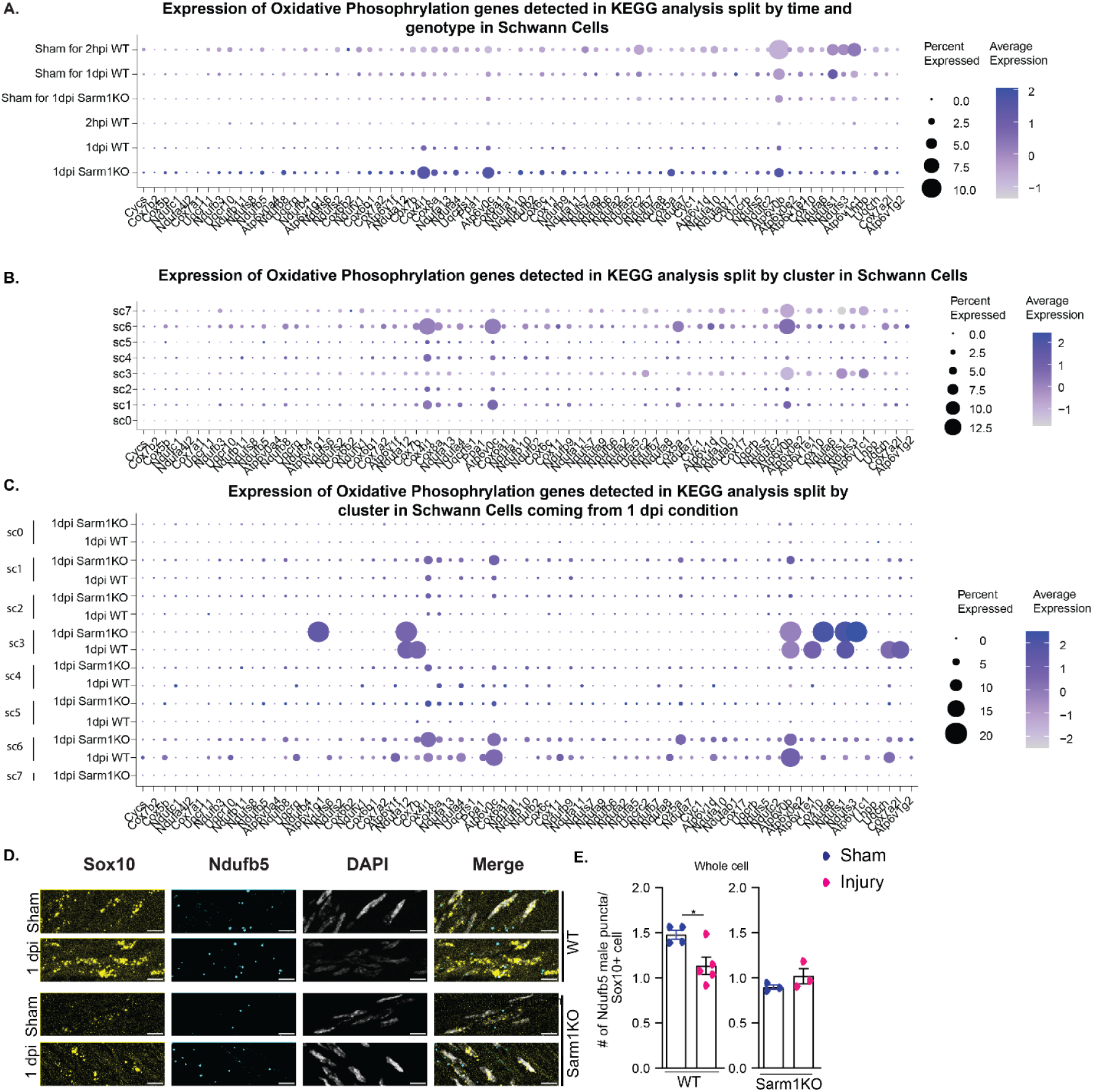
Supplemental information on all SCs. **A-C.** Dot plots showing expression of mitochondrial respiration genes (identified in Figure 3G) across all SCs, split by genotype and condition (Sham, 2hpi, 1dpi) (A), by cluster (B), and by cluster, condition, and genotype simultaneously (C). **D.** Representative images of RNA in situ hybridization for Ndufb5 (mitochondrial respiration gene) and Sox10 (SC marker) in WT and Sarm1KO sciatic nerves under sham conditions and 1 day post-injury. Scale bar = 10 µm. **E.** Quantification of Ndufb5 transcript numbers in Sox10-positive cells from WT and Sarm1KO sciatic nerves under sham and 1 day post-injury conditions. Data presented as mean ± SEM, p-value * = 0.0213.

**Supplemental Figure 6:**
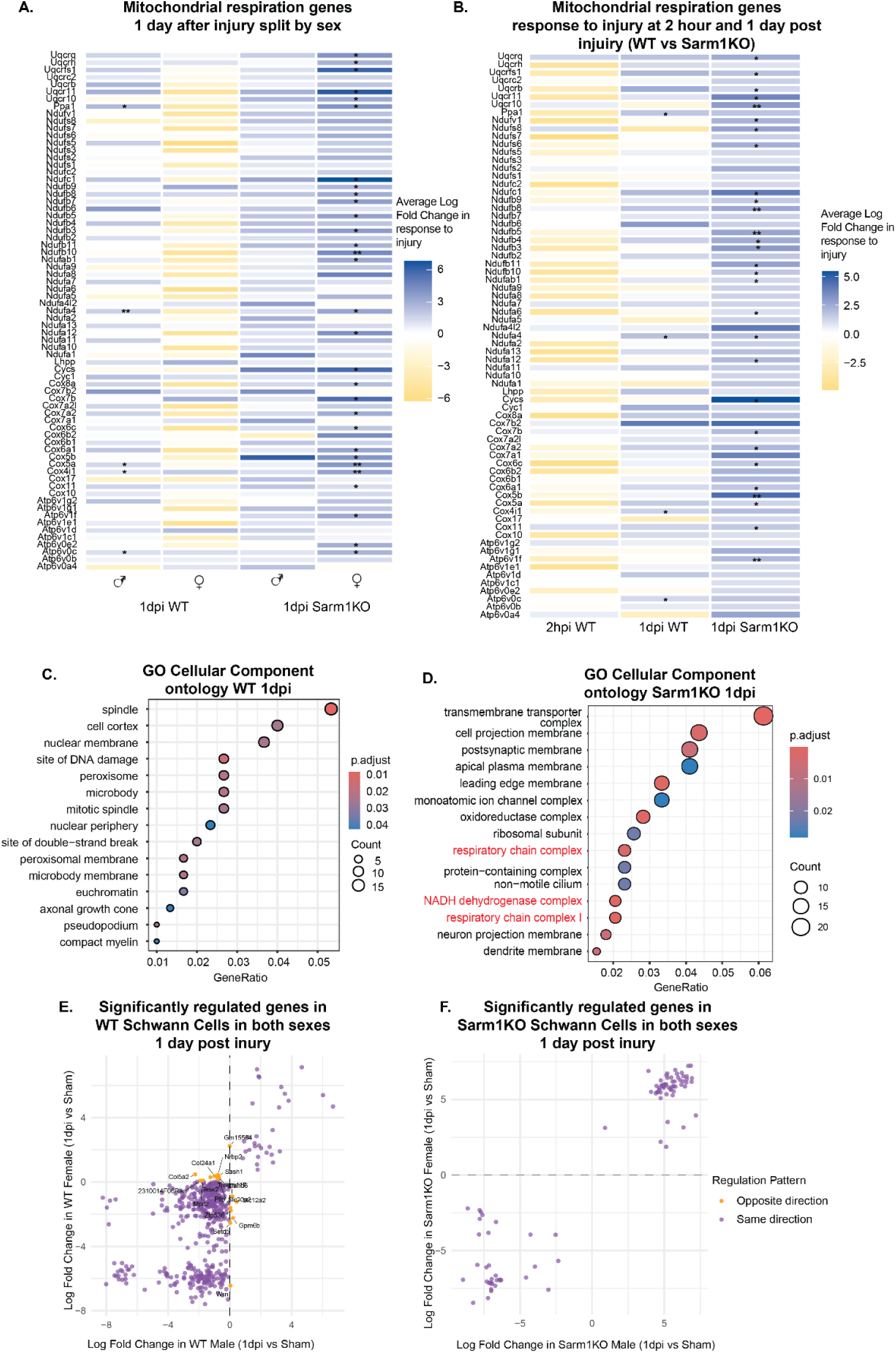
Supplemental information regarding sex differences and additional GO ontology analysis of mitochondrial respiration genes 1 day after injury. **A.** Average log fold change of mitochondrial respiration genes (identified in Figure 3G) 1 day after injury, split by sex. **B.** Expression of mitochondrial respiration genes (identified in Figure 3G) at 2 hours and 1 day after injury, split by genotype (WT vs Sarm1KO). **C-D.** GO cellular component ontology for WT (C) and Sarm1KO (D) SCs 1 day after injury. **E-F.** X-Y scatter plots showing gene expression log fold changes 1 day after injury, compared between sexes in WT SCs (E) and Sarm1KO SCs (F). In each plot, the x-axis represents gene changes in female SCs and the y-axis represents gene changes in male SCs. Only genes with an adjusted p-value < 0.05 (based on pseudobulk analysis for each sex) are shown. Purple highlights indicate genes regulated concordantly (both upregulated or both downregulated after injury in both sexes), while orange highlights indicate genes regulated discordantly (upregulated in one sex but downregulated in the other).

**Supplemental Figure 7:**
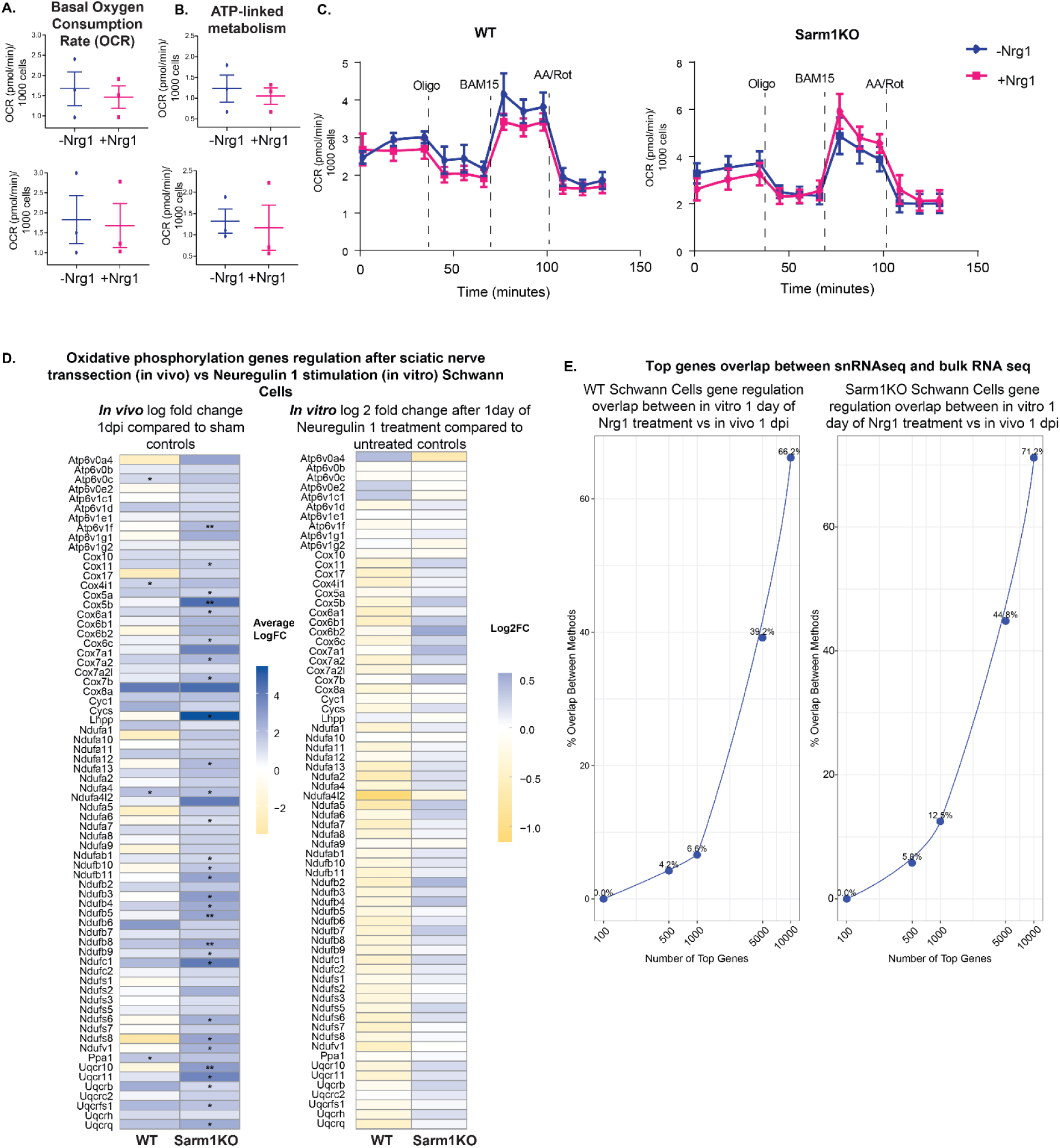
Oxygen consumption rate measurements and transcriptional responses in SCs. **A-B.** Oxygen consumption rate (OCR) measurements in primary mouse SCs from WT or Sarm1KO backgrounds using Seahorse XFe96 analyzer: (A) Basal OCR. (B) ATP-linked OCR. Each data point represents the mean of technical replicates from one mouse litter. **C.** Representative traces showing oxygen consumption rates (OCR) in WT (left) and Sarm1KO (right) SCs during mitochondrial stress test. Cells were treated with or without Neuregulin1 (Nrg1) for 24 hours to mimic injury conditions. OCR was measured at baseline and following sequential addition of oligomycin (Oligo), BAM15, and a combination of antimycin A and rotenone (AA/Rot) as indicated by vertical dashed lines. Data points represent mean ± SEM from technical replicates. Blue lines: control (-Nrg1); Pink lines: Nrg1-treated (+Nrg1). **D.** Comparison of mitochondrial respiration gene expression changes in SCs: snRNA-seq *in vivo* response to sciatic nerve transection injury 1 day post-injury versus *in vitro* bulk RNA-seq response to Nrg1 after 1 day of stimulation treatment. Log fold change is shown relative to sham nerves (*in vivo*) and untreated SCs (*in vitro*), p-value *<0.05, **<0.01, ***<0.001. **E.**Plots showing percentage overlap of top differentially expressed genes between snRNAseq of Schwann cells at 1 dpi and bulk RNAseq of Schwann cells after 24 hours of Nrg1 stimulation, for WT (left) and Sarm1KO (right) conditions. Overlap was calculated for the top 100, 500, 1,000, 5,000, and 10,000 differentially expressed genes, stratified by direction of regulation (upregulated or downregulated) in response to injury or Nrg1 stimulation.

## Table captions

Supplementary Table 1: Differential Gene expression analysis of each cluster detected in snRNAseq analysis

Supplementary Table 2: Differential Gene expression analysis of each cluster of Schwann Cells detected in snRNAseq analysis

Supplementary Table 3: Differential Gene expression analysis of each cluster of sham Schwann Cells detected in snRNAseq analysis

Supplementary Table 4: Pseudobulk analysis of differentially expressed genes between Wild type and Sarm1KO sham Schwann Cells

Supplementary Table 5: Differential Gene expression analysis of each cluster of 2 hours post injury (2hpi) and 1 day post injury (1dpi) Schwann Cells detected in snRNAseq analysis

Supplementary Table 6: Pseudobulk analysis of differentially expressed genes between Wild type sham and 1 day post injury Schwann Cells split by genotype (WT and Sarm1KO)

Supplementary Table 7: Pseudobulk analysis of differentially expressed genes between Wild type sham and 1 day post injury Schwann Cells split by genotype (WT and Sarm1KO) and sex

Supplementary Table 8: Pseudobulk analysis of differentially expressed genes between Wild type sham and 1 day post injury (1dpi) and sham and 2 hours post injury (2hpi) Schwann Cells split by genotype (WT and Sarm1KO)

**Table 9:**
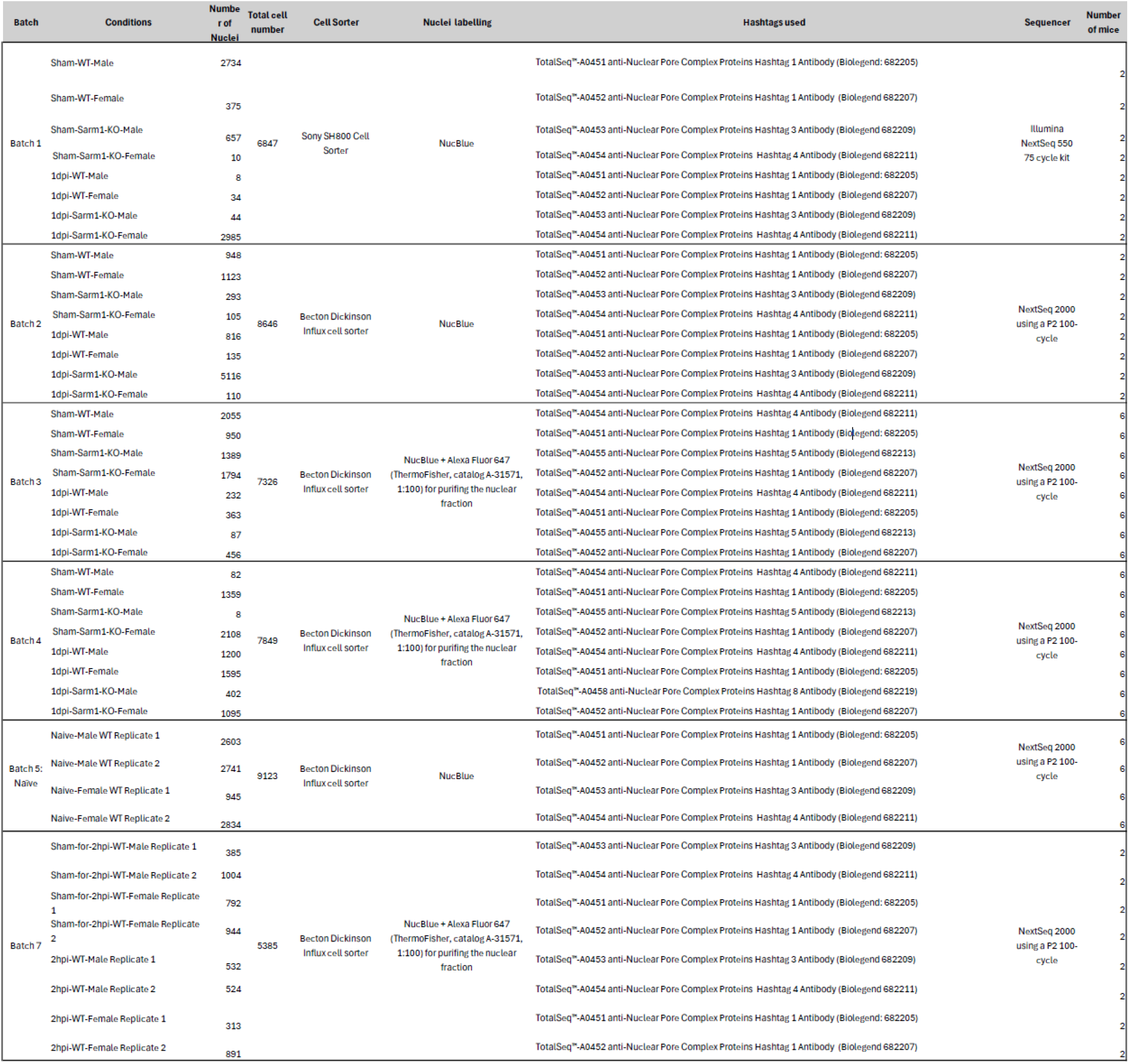
Overview of snRNAseq runs.

**Table 10:** Overview of Drosophila melanogaster experimental crosses.

| Purpose | Parent 1 | Parent 2 | Experimental progeny | Note |
| --- | --- | --- | --- | --- |
| Test dSarm in glia | w,hsFLP,FRT19A;Vglut-QF2,aseFlp2d/CyO; repoGal4, tubGal180t5,ascFLP/TM3sb,e | w,tubQSFRT19A;;UAS-LuciferaseRNAi/TM3sb,e | w,hsFLP,FRT19A/w,tubQSFRT19A;Vglut-QF2,aseFlp2d/+;UAS-LuciferaseRNAi/repoGal4, tubGal180t5,ascFLP | Take females only |
|  | w,hsFLP,FRT19A;Vglut-QF2,aseFlp2d/CyO; repoGal4, tubGal180t5,ascFLP/TM3sb,e | w,tubQS, FRT19A;UAS-RNAi dSarm#2/CyODgv | w,hsFLP,FRT19A/w,tubQSFRT19A;Vglut-QF2,aseFlp2d/dSarm#2;repo Gal4, tubGal180t5,ascFLP/+ | Take females only |
| Test dSarm RNAis in neurons | w;dpr1-Gal4, 5xUAS-Gal4, UAS-CD8-GFP, UAS-Dcr2/CyO; Dr/TM3 | w;+/CyODgv;UAS-RNAiLuciferase/TM6BTb, Hu,e | w;dpr1-Gal4, 5xUAS-Gal4, UAS-CD8-GFP, UAS-Dcr2/+;UAS-Luciferase/Dr |  |
|  | w;dpr1-Gal4, 5xUAS-Gal4, UAS-CD8-GFP, UAS-Dcr2/CyO; Dr/TM3 | w;UAS-RNAi dSarm#1 | UAS-CD8-GFP, UAS-Dcr2/w;UAS-RNAi dSarm#1; Dr/+ |  |
|  | w;dpr1-Gal4, 5xUAS-Gal4, UAS-CD8-GFP, UAS-Dcr2/CyO; Dr/TM3 | w;UAS-RNAi dSarm#2 | UAS-CD8-GFP, UAS-Dcr2/w;UAS-RNAi dSarm#2; Dr/+ |  |

